# Frontal cortex activity underlying the production of diverse vocal signals during social communication in marmoset monkeys

**DOI:** 10.1101/2021.08.30.458300

**Authors:** Lingyun Zhao, Xiaoqin Wang

## Abstract

Vocal communication is essential for social behaviors in humans and many non-human primates. While the frontal cortex has been shown to play a crucial role in human speech production, its role in vocal production in non-human primates has long been questioned. Recent studies have shown activation in single neurons in the monkey frontal cortex during vocal production in relatively isolated environment. However, little is known about how the frontal cortex is engaged in vocal production in ethologically relevant social context, where different types of vocal signals are produced for various communication purposes. Here we studied single neuron activities and local field potentials (LFP) and in the frontal cortex of marmoset monkeys while the animal engaged in vocal exchanges with other conspecifics in a social environment. Marmosets most frequently produced four types of vocalizations with distinct acoustic structures, three of which were typically not produced in isolation. We found that both single neuron activities and LFP were modulated by the production of each of the four call types. Moreover, the neural modulations in the frontal cortex showed distinct patterns for different call types, suggesting a representation of vocal signal features. In addition, we found that theta-band LFP oscillations were phase-locked to the phrases of twitter calls, which indicates the coordination of temporal structures of vocalizations. Our results suggested important functions of the marmoset frontal cortex in supporting the production of diverse vocalizations in vocal communication.

## Introduction

Many non-human primate species use vocalizations during their social communication. Field studies and experiments from the lab have shown that these vocalizations provide important functions for their social behaviors (1). Evidence has been found that the control and use of the vocalizations resemble some of the rudimentary features in human speech and communication (1–3). However, the neural mechanism underlying vocal production and communication remains largely unknown. Studies in humans have identified that the production of emotional vocalizations (e.g. cry and laughter) was processed in structures on the medical side of the brain, including the anterior cingulate cortex and periaqueductal gray (PAG) (4). In contrast, the production of communicative signals (e.g. speech) was found to be processed in structures on the lateral side of the brain, including Broca’s area and multiple premotor and motor regions in the frontal cortex (4). These lateral structures were especially important for the timing (5, 6) and the generation of distinct acoustic structures (7–10) in speech. A large body of previous studies in non-human primates focused on vocalizations that were induced by operant conditioning or electrical stimulation (11, 12). The brain structures identified in these studies were largely on the medical side, such as the anterior cingulate cortex and PAG, similar to what is responsible for emotional vocalizations in humans. The prefrontal cortex and the premotor cortex were found to be active for conditioned vocalizations but not spontaneous vocal production (13, 14). It is a long-standing question whether these lateral structures in the frontal cortex are involved in natural vocal production in monkeys.

There has been growing evidence for the marmoset (*Callithrix jacchus*), a New World monkey species, to have a higher level of vocal plasticity and flexibility than previously thought in non-human primates (15). For example, the vocal development in juvenile marmosets was influenced by social feedback from parents through vocal interactions (16–18). Adult marmosets demonstrated flexibility and voluntary control in some aspects of vocal production, including the initiation time (2), acoustic structures (19–21) and vocal turn-taking (3, 22). Vocal structures were also found to be modulated by social contexts (23–25). These findings in behavioral studies draw attention to the frontal cortex for its potential involvement in vocal control.

To study the correlation between the frontal cortex and vocal communication in non-human primates, it is important to engage the subjects in social contexts where the ethological function of their vocalizations was preserved. Using an antiphonal calling paradigm (26, 27), recent studies have found neural activities in the premotor and prefrontal cortices of the marmoset monkeys were modulated when the marmosets produced phee calls in exchange with another conspecific (28–30). Although these data provided initial evidence, they had limitations as phee calls were produced mostly in isolation. Marmosets produce more than 15 types of vocalizations during natural social communication (31–33). It is unclear whether the frontal cortex plays a role in the generation of these social communication calls. One hypothesis would be that the frontal cortex does not show any activity to vocal production other than phee calls, which may indicate that vocal communication is largely controlled by medial and subcortical structures. However, the frontal cortex may be active for a subset of, or even all call types, if these call types are interfaced with some degrees of cognitive process or voluntary control. If this is true, we would further ask what specific function the frontal cortex has regarding the control of these calls. It could be a timing signal for call initiation, in which case the activity may not distinguish different types of calls but simply drive subcortical structures to create the vocal structure for particular call types. Alternatively, the activity can be a motor control-related signal that is correlated with individual call types.

To address these questions, we performed our current study in the marmoset breeding colony, in which dozens of marmosets were housed together with their social communications maintained in a naturalistic way. Individual marmosets generated all types of calls within the vocal repertoire and exchanged them with other marmosets in the room. To enable experiments in such an environment, we developed new techniques to overcome challenges from noise interference in acoustic and wireless neural recordings. We hypothesize that the frontal cortex is active during the vocal production of social communication calls and the neural activity represents the particular call type being generated. We focused on four major types of calls used in marmoset communication, each of which has distinct acoustic structures. We found local-field potentials and single neurons were both modulated by each of these call types. Further analysis showed that these activities distinguish the call types being produced and provide temporal information for the vocal structures within a call. These results have important implications on the functional role of the marmoset frontal cortex in vocal production and communications.

## Results

We recorded local field potentials (LFP) and single neuron activities in the frontal cortex of the left hemispheres from two marmosets while they were freely roaming in a cage in the marmoset colony and engaged in social communications with conspecifics housed in the same room (Fig. 1, see Materials and Methods, Supplementary Methods, SI Appendix). One of the subjects was implanted with a 32-channel electrode array (Subject M93A, brown rectangle in Fig. 1C) and the other subject was implanted with a 16-channel electrode array (Subject M9606, cyan square Fig. 1C). In each session, a wireless headstage was coupled to the electrode array and the experimental subject engaged in its natural behaviors, including eating, drinking, playing and vocalizing without being interfered with by the recording setup. There was no other marmoset housed in the same cage with the subject, but the subject was able to see and hear other marmosets in the neighboring cages and other parts of the room, including the subject’s family members. The subject generated vocalizations of various call types within the entire vocal repertoire of marmosets (31). To ensure reliable recordings, we developed a new apparatus and routine for wireless neural recording based on a previously built system in the lab (Fig. 1A, B) (34). We developed a parabolic microphone system to identify vocalizations of the experimental subject from the noisy background (Fig. 1D-F, see Supplementary Methods, SI Appendix). Neural signals and vocalizations were continuously recorded during the experimental session for two to five hours each day. Only the four major types of vocalizations (Fig. 2A-D) were included in the analysis (31).

**Fig. 1.**
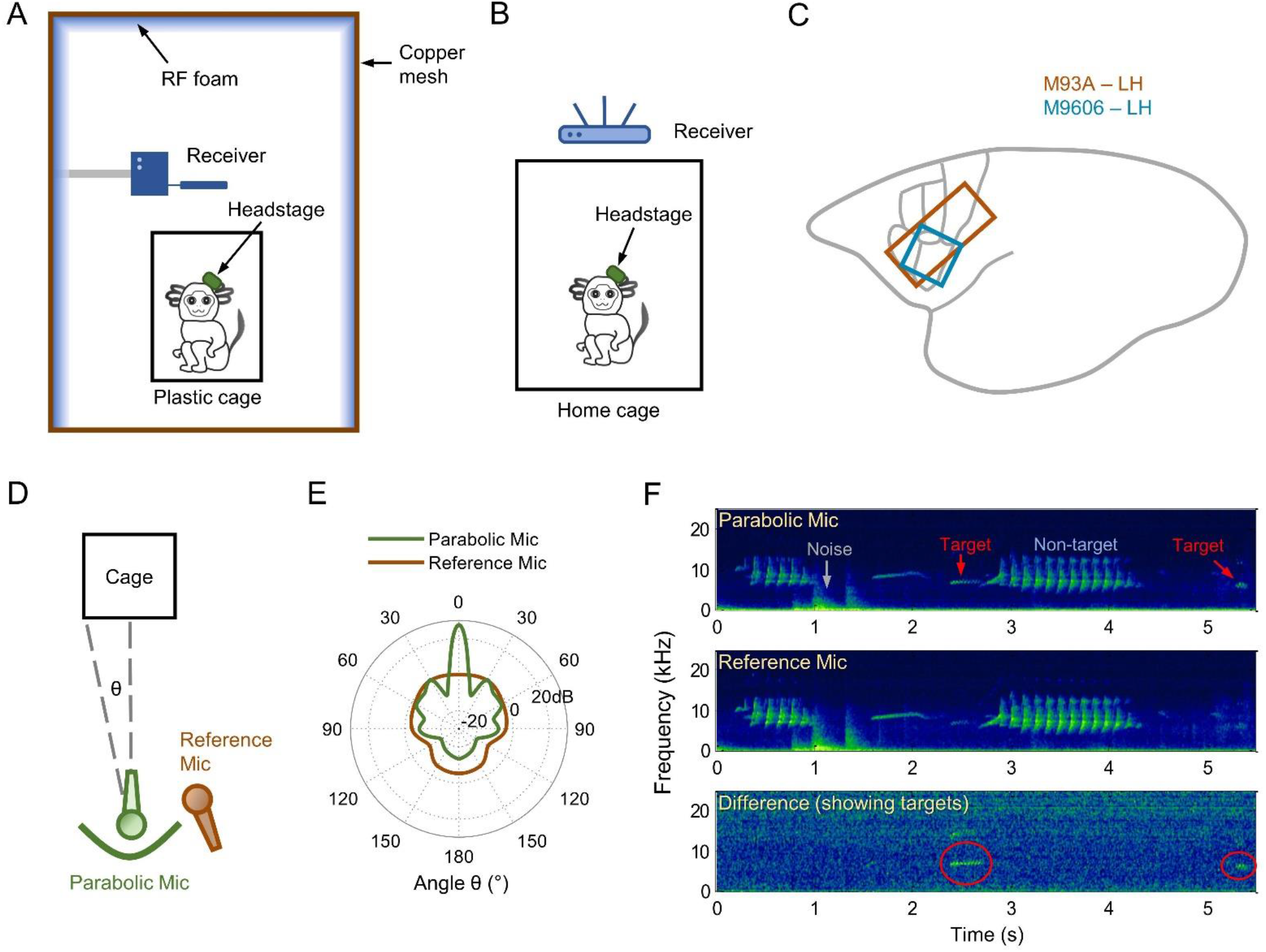
Experimental setup in the marmoset colony. (**A**) Schematic of the wireless neural recording setup (side view) using the analog system (for Subject M9606). The subject and the wireless receiver were enclosed by a shielded booth to minimize interference during wireless signal transmission. The shielded booth was constructed with copper mesh walls and lined with RF absorption foam. (**B**) Schematic of the wireless neural recording setup (side view) using the digital system (for Subject M93A). (**C**) Illustration of the areas (colored rectangles) covered by electrode arrays within the marmoset cortex for the two subjects. Brown: M93A, 32 channels. Cyan: M9606, 16 channels. LH: left hemisphere. The gray lines near the rectangles outline the borders of several Brodmann areas covered by the arrays (see Fig. 2I-P). (**D**) Schematic of the directed acoustic recording setup (top view). Microphones were placed in front of the cage where the subject (target) was housed. Green: parabolic mic. The green curve illustrates the parabolic reflector. Brown: reference mic. (**E**) Pick-up pattern (gain at different angles) of the two types of microphones at 8kHz. The gain of the parabolic mic peaks at the zero-degree angle (front-facing direction) and quickly drops as the angle increases (side directions). Within a certain angle range (a narrow range near front-facing direction), the gain of the parabolic mic is higher than the reference mic. (**F**) An example recording clip (spectrograms) from the parabolic-reference mic pair in the colony. Top: channel from the parabolic mic. Background noise, non-target vocalizations and the target vocalizations from the subject were recorded. The target vocalization has an enhanced intensity. Middle: channel from the reference mic. Background noise, non-target vocalizations and weaker target vocalizations were recorded. The intensity of background noise and non-target vocalizations in the parabolic channel is similar to that in the reference channel, whereas the intensity of the target vocalizations is much stronger in the parabolic channel. Bottom: the difference between the parabolic channel and the reference channel. Background noise and non-target vocalizations cancel out. Only target vocalizations remain.

**Fig. 2.**
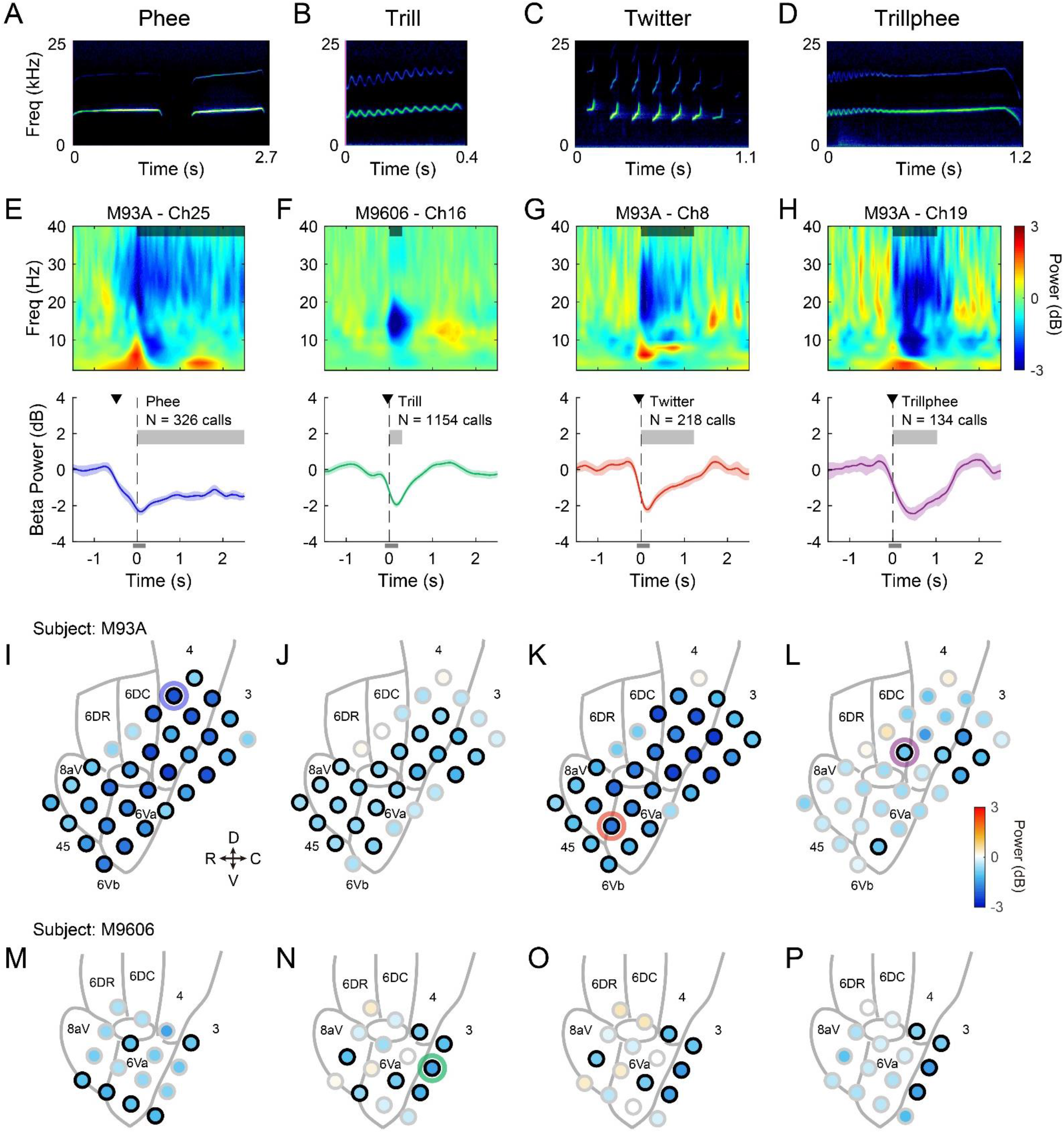
Suppression in the beta-band local field potential (LFP) for four types of marmoset calls: phee (**A,E,I,M**), trill (**B,F,J,N**), twitter (**C,G,K,O**) and trillphee (**D,H,L,P**) (**A**) Spectrogram of an example phee call from one marmoset (Subject M93A, with two syllables). (**B-D**) Same format as **A**, spectrograms of trill, twitter and trillphee calls. (**E**) LFP from one example recording site showing suppression in beta-band (12-30Hz) when the subject vocalized phee calls. The location of the site is indicated by a blue circle in **I**. Top: time-frequency representation of LFP. The color scale is shown next to **H**. The shaded bar near the top of the panel illustrates the average call duration (truncated at the axis limit). LFP signals were aligned to the vocal onset (time zero) for each call and the time-frequency representations were averaged across calls. The subject ID and the channel number for the recording site are indicated above the panel. Bottom: LFP power in the beta band (mean ± SEM) relative to that in the baseline window before vocal onset. Vertical dashed line: time zero. Thick gray bar: average call duration (truncated at the axis limit). N indicates the total number of calls included. The black triangle indicates the suppression start time, i.e. the earliest time at which LFP power showed significant suppression (see **Fig. 3C**). The thin gray bar below the axis indicates the analysis window near vocal onset used to quantify the modulation of beta-band power in **I-P**. (**F-H**) Same format as **E**, for the three other call types. (**I**) Spatial distribution of beta-band modulation across all recording sites for phee calls in Subject M93A. Each circle indicates a recording site. The color in the circle indicates the LFP power in the analysis window shown in **E** (a thin bar below the bottom panel) relative to that in the baseline window before vocal onset. The color scale is shown next to **L**. A black border on a circle indicates significant modulation. The blue ring indicates the location of the example site in **E**. Gray lines in the background outline the border of Brodmann areas according to the marmoset brain atlas with their names labeled. A small area near the center without a label is area 8C. R: rostral; C: caudal; D: dorsal; V: ventral. (**J-L**) Same format as **I**, for the other three call types in Subject M93A. (**M-P**) Same format as **I**, for Subject M9606.

### Beta-band suppression for the production of social communication calls

We analyzed modulations in LFP signals when marmosets produced vocalizations during social communication. LFP activity in the beta-band (12-30Hz) has long been observed in the frontal motor areas during resting and motor movements in humans and animals (35, 36). Beta-band suppression was found during motor preparation and execution of voluntary movements and was thought to be related to the synchronization of neural activities (35–38). We first ask whether there is any modulation in the motor areas of the marmoset frontal cortex that is comparable to what was often seen for voluntary movement (Fig. 2).

The four major call types for social communication have distinct spectrotemporal features in the acoustic structure (Fig. 2A-D). Three of the four call types are narrowband (phee, trill and trillphee) whereas the other one is wideband (twitter). A phee call is a long and loud call with mainly linear frequency modulations. It is usually composed of one or more syllables (Fig. 2A). A trill call is relatively short and weak in its intensity, with sinusoidal frequency modulations (Fig. 2B). A twitter call is composed of multiple short syllables of sharply rising frequency modulations (Fig. 2C). A trillphee call starts with sinusoidal frequency modulations like in a trill and then dampens into linear frequency modulations like in a phee (Fig. 2D). Figure 2E top panel shows the time-frequency representation of LFP recorded from one example site in the motor cortex for phee calls (shown for up to 2.5 sec after vocal onset, see Fig. 2I blue ring for the location of the site). The shaded bar near the top indicates the average duration of phees (truncated at 2.5 sec). The strongest modulation was observed in the beta band. To quantify such modulation, we filtered the LFP signals for the beta band (12-30Hz) and compared the averaged power at different time points (Fig. 2E, bottom) to that in the baseline window ([−3,-1] sec). Beta-band LFP showed a decrease in power before and during the production of phee calls, consistent with the beta-band suppression in motor areas in the previous literature. The suppression started before the vocal onset and had the largest effect around vocal onset. The earliest time at which the beta power showed significant suppression (i.e. suppression start time) for this site is −0.485 sec (Fig. 2E, black triangle in the bottom panel, p < 0.05, signed-rank test). Similar modulation to the beta-band LFP was observed for trill, twitter and trillphee calls from various recording sites (Fig. 2F-H). The beta power started to decrease near the vocal onset (−0.045 sec for trills, −0.055 sec for twitters, −0.010 sec for trillphees, p < 0.05, signed-rank test) and had its largest decrease near the beginning of or in the early phase of the calls. Therefore, beta-band suppression was observed for the production of all four major types of marmoset vocalizations. To compare the beta-band suppression across cortical regions, we calculated the beta power for each site within an analysis window near vocal onset ([−0.1, 0.2] sec, Fig. 2E-H, gray bars below the bottom panels) and illustrate the size and significance of the suppression effect across cortical regions (Fig. 2I-P, black border for significant suppression, p < 0.05, signed-rank test). For phee, trill and twitter calls, the sites with beta-band suppression were not restricted to a particular region but seemed to scatter across multiple regions in the frontal cortex. The sites with suppression for trillphee calls seemed to be located in the more caudal regions (Fig. 2L, P).

We then ask whether the modulation in the beta-band LFP simply indicates the generation of any vocalization or depends on call types. We compared the beta-band suppression between the four different call types at each recording site. At one example recording site, the suppression for four call types has different temporal profiles (Fig. 3A, left). Within the analysis window near vocal onset, the average power showed a significant difference between several pairs of call types (Fig. 3A, right, p < 0.05 for each pair, One-way ANOVA, *post-hoc* analysis with Bonferroni corrections). Phee calls induced the largest suppression and trill induced the smallest. Trillphees have their acoustic structure combining features of both trill and phee (Fig. 2D) and the size of suppression for it is between that for phees and trills. Another example recording site showed suppression with different amplitude and temporal profiles for the four call types (Fig. 3B). In total, 28 out of 32 sites in Subject M93A and 8 out of 16 sites in Subject M9606 showed differences in beta power in the analysis window for different call types.

**Fig. 3.**
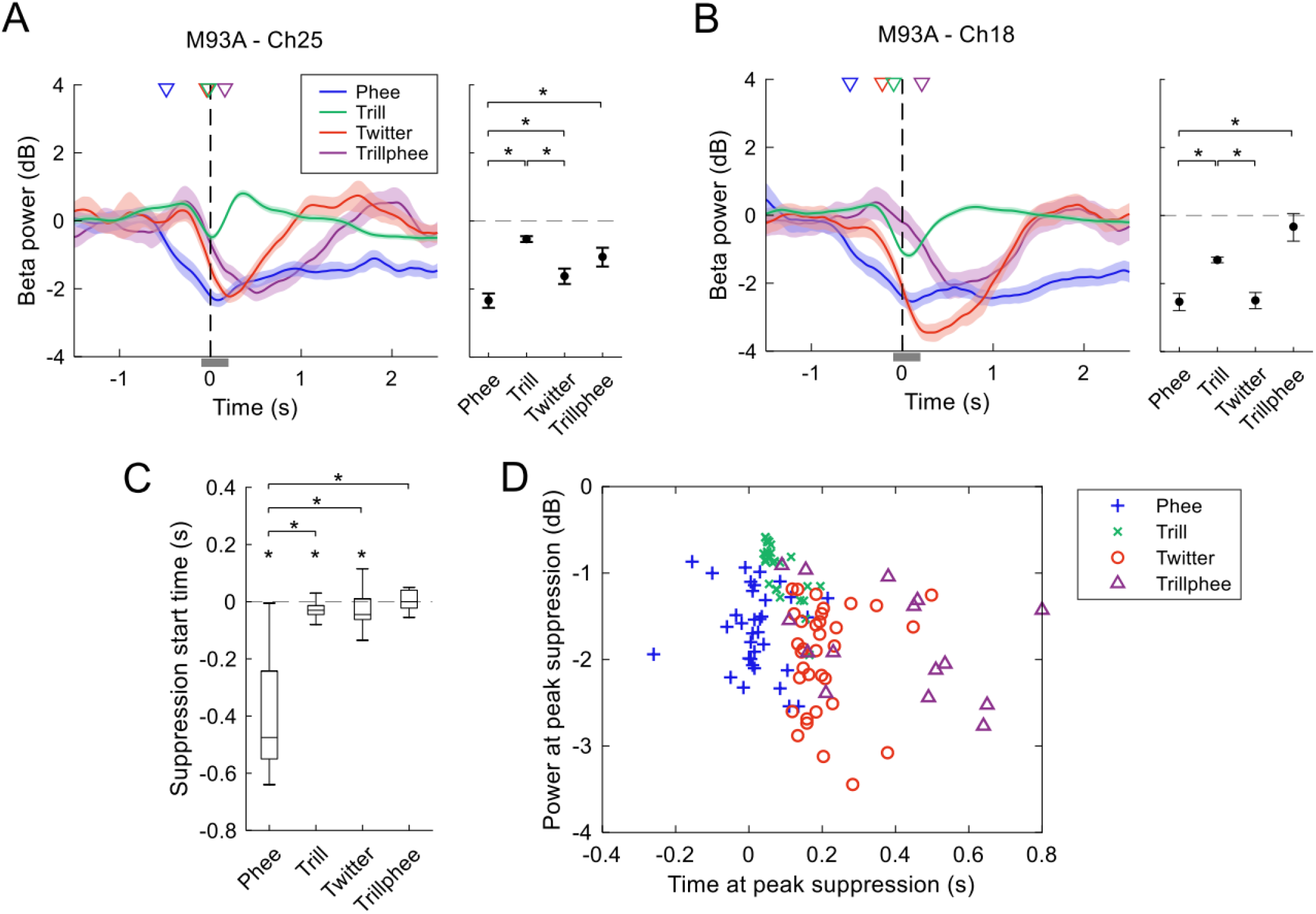
Comparison of the beta-band suppression between four call types. (**A**) Left: LFP power in the beta band (mean ± SEM) from one example recording site for four call types. Blue: phee; green: trill; red: twitter; purple: trillphee. The LFP power is relative to that in the baseline window. The subject ID and channel number are labeled on top. The colored triangles indicate the suppression start time, i.e. the earliest time at which LFP power showed significant suppression. The thin gray bar indicates the analysis window to quantify LFP power (see the right panel). Right: LFP power in the beta band (mean ± SEM) in the analysis window shown in the left panel. An asterisk indicates a significant difference between two call types. (**B**) Same format as **A**, for another recording site. (**C**) Comparison of the suppression start time between call types (Tukey boxplot). Data are from all recording sites showing significant suppression. Medians: horizontal lines inside the boxes. First and third quartiles: lower and upper borders of the boxes. Inner fences: whiskers outside of the boxes. Outliers are not plotted. An asterisk above a whisker indicates that the median is significantly lower than zero. An asterisk above a bracket indicates a significant difference between two call types. (**D**) Scatter plot of the time and power at peak suppression for four call types. Each data point is from a recording site showing significant suppression for the call type indicated by color. Blue plus: phee; green cross: trill; red circle: twitter; purple triangle: trillphee. One data point with the time at peak suppression greater than 0.8 sec is plotted on the border of the axes.

To quantify the difference shown in the beta power (Fig. 3A, B), we pooled the suppression start time from the population of sites that showed significant suppression to each call type (see Materials and Methods), we found that the median suppression start time is below zero for phee, trill and twitter calls (Fig. 3C, p < 0.05, signed-rank test), i.e. the suppression started before the vocal onset. Interestingly, the suppression for phee calls started significantly earlier than the other three call types (Fig. 3C, p < 0.05 Kruskal-Wallis test, *post-hoc* analysis with Bonferroni corrections). We further summarize the difference in the temporal profile in the beta power (Fig. 3A, B) for population data. For each call type, we plot the power and time at peak suppression for the sites that showed significant suppression (Fig. 3D). The four groups of data points in the scatter plot do not overlap entirely. Instead, a subset in each group occupies a separate region in the coordinate plane. This shows that, for each call type, there is a subpopulation of sites that shows unique activities in the beta-band LFP for that call type. It is interesting that the size of the suppression induced by trillphees spans in a similar range as that induced by phee calls (Fig. 3D, power at peak suppression). However, the time when suppression reaches its peak for trillphee is usually later than that for phee (Fig. 3D, time at peak suppression). A possible explanation for this may come from the fact that a trillphee starts with a trill-like first component followed by a phee-like second component. If the modulation in the first component follows that for trills, then it will start with a small suppression. If the modulation in the second component follows that for phees, then the suppression will increase in the middle of a trillphee call. Combining the modulation in the two components will result in the time at peak suppression being later than that for phee calls, which is in the middle of trillphees. With the above observations, our data suggest that the beta-band suppression not only reflects the vocal production during social communication but also represents call type information both from individual sites and as a population.

### Theta-band activation and phase lock with twitter syllables

Besides beta-band suppression, we observed modulations in other frequency bands in LFP. Interestingly, a subset of sites showed activation in the theta-band (4-8Hz). An example recording site showed an increase in LFP power in the theta band for the production of twitter calls (Fig. 4A, left panels). The theta-band activation started before the vocal onset of twitter calls (Fig. 4A, black triangle in the bottom left panel), peaked at the beginning of calls and lasted for the duration of production. Another example site showed similar activation for twitters (Fig. 4A, right panels). This activation in the theta band is also found in other call types, although with a smaller number of sites. An example of theta-band activation induced by trill calls is shown in Figure 4B, with a smaller power increase compared to the examples for twitters shown in Figure 4A. When averaging the theta power within the call duration for each site, we found significant activation in two sites for phees, five sites for trills, fifteen sites for twitters and two sites for trillphees in Subject M93A, plus two sites for twitters in Subject M9606. If we include sites with any activation before or during the vocalizations and summarize the activation start time (i.e. the earliest time at which LFP power showed significant activation, see Materials and Methods), we found that the median of activation start time is significantly below zero for phee, trill and twitter calls (Fig. 4C, p < 0.05, signed-rank test). There is no significant difference in the median between the four call types (Fig. 4C, p > 0.05, Kruskal-Wallis test).

**Fig. 4.**
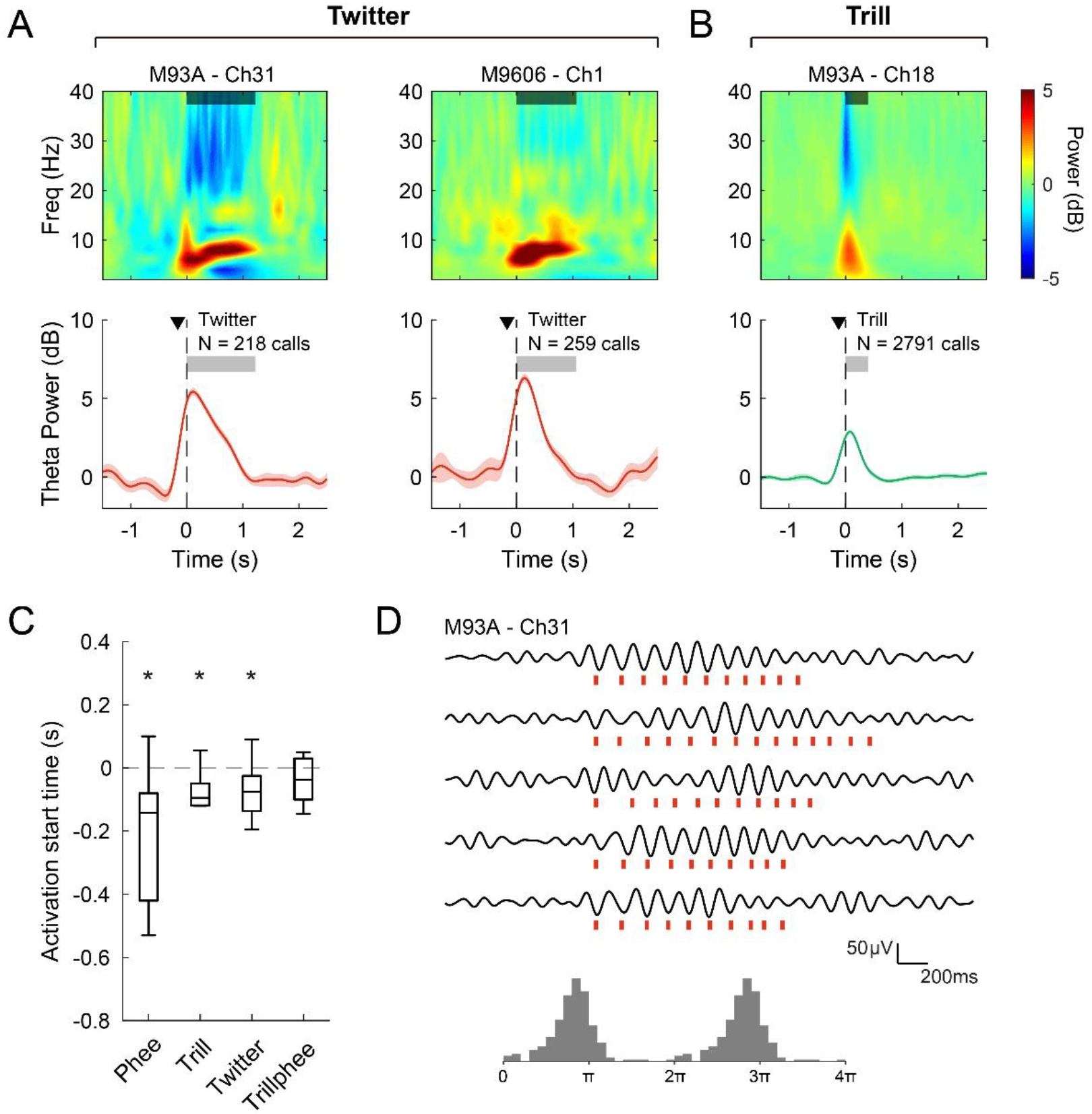
Activation in the theta-band LFP. (**A**) Left: same format as **Fig. 2E**, LFP from one example recording site (in Subject M93A) showing activation in theta band (4-8Hz) when the subject vocalized twitter calls. The color scale is shown next to **B**. The black triangles in the bottom panel indicate the activation start time, i.e. the earliest time at which LFP power showed significant activation (see **C**). Right: same format as the left panel, for an example site from the other subject (M9606). (**B**) Same format as the left panel of **A**, activation in the theta-band LFP for trill calls. (**C**) Same format as **Fig. 3C**, comparison of the activation start time between call types (Tukey boxplot). Data are from all recording sites showing significant activation. An asterisk above a whisker indicates that the median is significantly lower than zero. There is no significant difference in the activation start time across the four types of calls. (**D**) Phase lock of the individual syllables of a twitter call to the theta-band LFP oscillations. Top: five example trials showing the theta-band LFP oscillations (black traces) from one recording site. The red bars below each trace indicate the onset time of each twitter syllable. Bottom: histogram of the phase of the theta-band LFP oscillations corresponding to the onset time of each twitter syllable from all twitter calls at this recording site.

It is noteworthy that a twitter call is composed of a series of fast-repeating short syllables (Fig. 2C). The repetition rate of the syllables is around 7Hz (31), which is within the frequency range of the theta band. We wonder whether the LFP activity in theta band is related to the repeated production of the twitter syllables. We therefore compared the LFP waveform in the theta band with the onset time of each twitter syllable (Fig. 4D). As shown in the example trials (Fig. 4D, top), LFP waveform showed increased amplitude during the production of twitters, as expected from the theta-band activation (Fig. 4A, left panels). Interestingly, the onset of twitter syllables occurred around a particular phase in the LFP oscillation – near the valley of each cycle. If the period of the oscillation cycle is longer, the interval between the syllables gets longer as well (Fig. 4D, second trace from top). Sometimes the syllable production skipped one cycle and the next syllable will still occur at a similar phase in the oscillation (Fig. 4D, third trace from top). These observations strongly suggested phase lock between the twitter syllables and the theta-band LFP oscillations in this site. Figure 4D bottom panel shows the distribution of the theta-band LFP phase from all twitter syllables in this recording site at their onset time. We used vector strength (VS) and Rayleigh statistics (p < 0.001 for Rayleigh > 13.8) to quantify the effect of phase lock. While vector strength characterizes how well each twitter syllable synchronizes with the theta-band LFP oscillations, Rayleigh statistics tests if the phase at twitter syllable onset has a uniform distribution (no phase-lock) or not. For this example site, the vector strength is 0.76 (Rayleigh: 239).

Across recording sites, we found that activation in the theta-band LFP power was more prominent towards the dorsal side of the arrays, near the primary motor cortex (Fig. 5A, B, average power within the duration of twitter calls). For Subject M93A, activation was found across several cortical regions, with the strongest effect in the primary motor cortex (Fig. 5A). For Subject M9606, activation was found in two sites at the border of the dorsal premotor cortex, Brodmann area 8C and primary motor cortex (Fig. 5B). In addition, two sites in Subject M93A in the ventral part of the premotor cortex showed suppression in theta power (Fig. 5A). For the phase-lock of twitter syllables, we found a majority of sites showed significant phase-lock (p < 0.001, Rayleigh > 13.8), except for one site in Subject M93A and one site in Subject M9606 (both in the ventral part of the array). In general, sites towards the dorsal part of the arrays showed larger vector strength, indicating a stronger phase lock between individual twitter syllables to the theta oscillations in LFP (Fig. 5C, D).

**Fig. 5.**
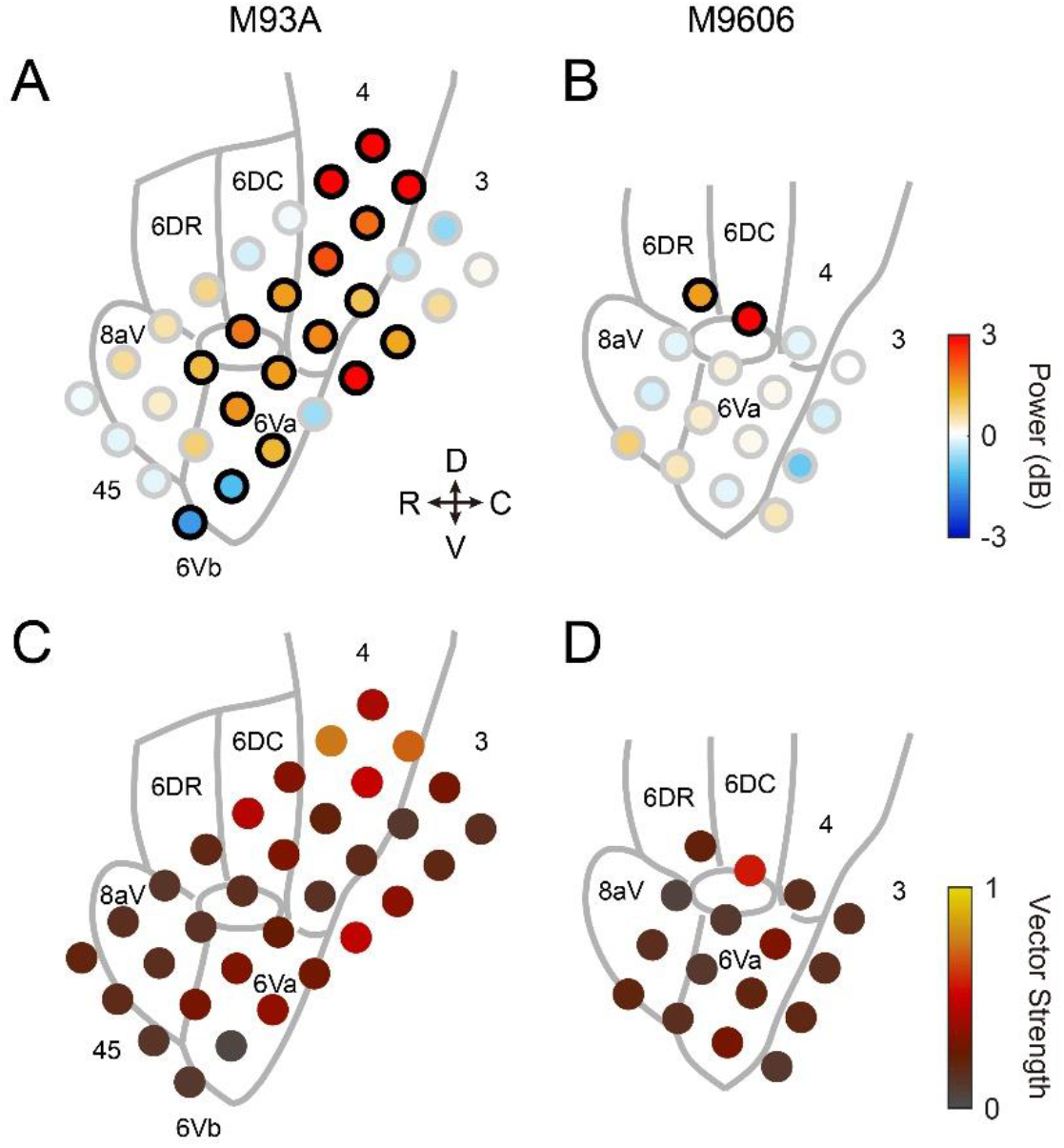
Spatial distribution of the theta-band modulation (**A, B**) and the phase lock (**C, D**) across all recording sites for twitter calls. (**A**) Same format as **Fig. 2I**, theta-band modulation for twitter calls (Subject M93A). The LFP power is calculated within an analysis window covering the duration of each twitter call. The color scale is shown next to **B**. A black border on a circle indicates significant modulation. (**B**) Same format as **A**, for Subject M9606. (**C**) Spatial distribution of the vector strength across all recording sites for twitter calls (Subject M93A). The color scale is shown next to **D**. (**D**) Same format as **C**, for Subject M9606.

### Single neuron activities modulated by social communication calls

Modulations to single neurons in the marmoset frontal cortex have been found in previous studies during vocal production of phee calls (28, 29). In our current experiment, we ask whether there is neuronal modulation during vocal production in the social communication context and if so, whether the modulation distinguishes different call types with distinct acoustic structures. Marmosets produced only a small number of phee calls in the colony and the number associated with each neuron was usually too small for calculating averaged firing rates. Therefore, we only included trill, twitter and trillphee calls in the analysis for single neurons. We consider a neuron as tested for one call type (one of trill, twitter or trillphee) if there were at least 15 calls of that type recorded from the neuron. In total, we obtained 151 neurons tested for at least one call type in our analysis (96 for Subject M9606; 55 for Subject M93A).

Figure 6 illustrates a few example neurons with firing rate modulated by the production of trill, twitter and trillphee calls. Spike timing is aligned to the vocal onset of each call. From the raw spiking activities, we observed large variations in the firing rate across trials in many neurons (e.g. Fig. 6C, top), likely due to the nature of free-roaming natural behavior. To reduce the variations across trials and reveal the modulation induced by vocalizations, we z-scored the firing rate for each trial and calculated the averaged activities (Fig. 6, bottom, see Materials and Methods). We used four analysis windows (early, pre-call, during-call and post-call windows) and compared the neural activities in each of these windows to that in the baseline window to determine whether a neuron is modulated by a particular call type (see Materials and Methods). For trill calls, one example neuron showed increased activities in the pre-call window (Fig. 6A). The activity started to increase early before the vocal production (more than one second before vocal onset) and peaked within half a second before vocal onset. Another example neuron showed decreased activities in the during-call window (Fig. 6B). For twitter calls, an example neuron showed increased activities in the post-call window (Fig. 6C). The activity first had a slight decrease near vocal onset and then increased and peaked right after the end of twitter calls. For trillphee calls, an example neuron showed increased activities in the during-call window (Fig. 6D). The total number of neurons modulated for each call type is summarized in Table 1. Overall, 31% (47/151) of neurons showed modulation to as least one of the three social communication call types. We then took a close look at when these modulations occurred relative to the four analysis windows. We calculated the proportion of neurons showing activation or suppression in each analysis window for each call type (Fig. 7A). Interestingly, trill calls induced activations at a very early time before the vocal onset (early window) in a subset of neurons (Fig. 7A, top). Twitter calls induced strong suppression during vocalizations and activations after a call was finished (Fig. 7A, middle). When we repeated the calculation with shuffled spike timing, which quantified any possible modulation effect by chance, the proportion of neurons in each window became very low (Fig. 7B). Therefore, the observations we had for the original data (Fig. 7A) did not arise from random fluctuation in spike activities.

**Fig. 6.**
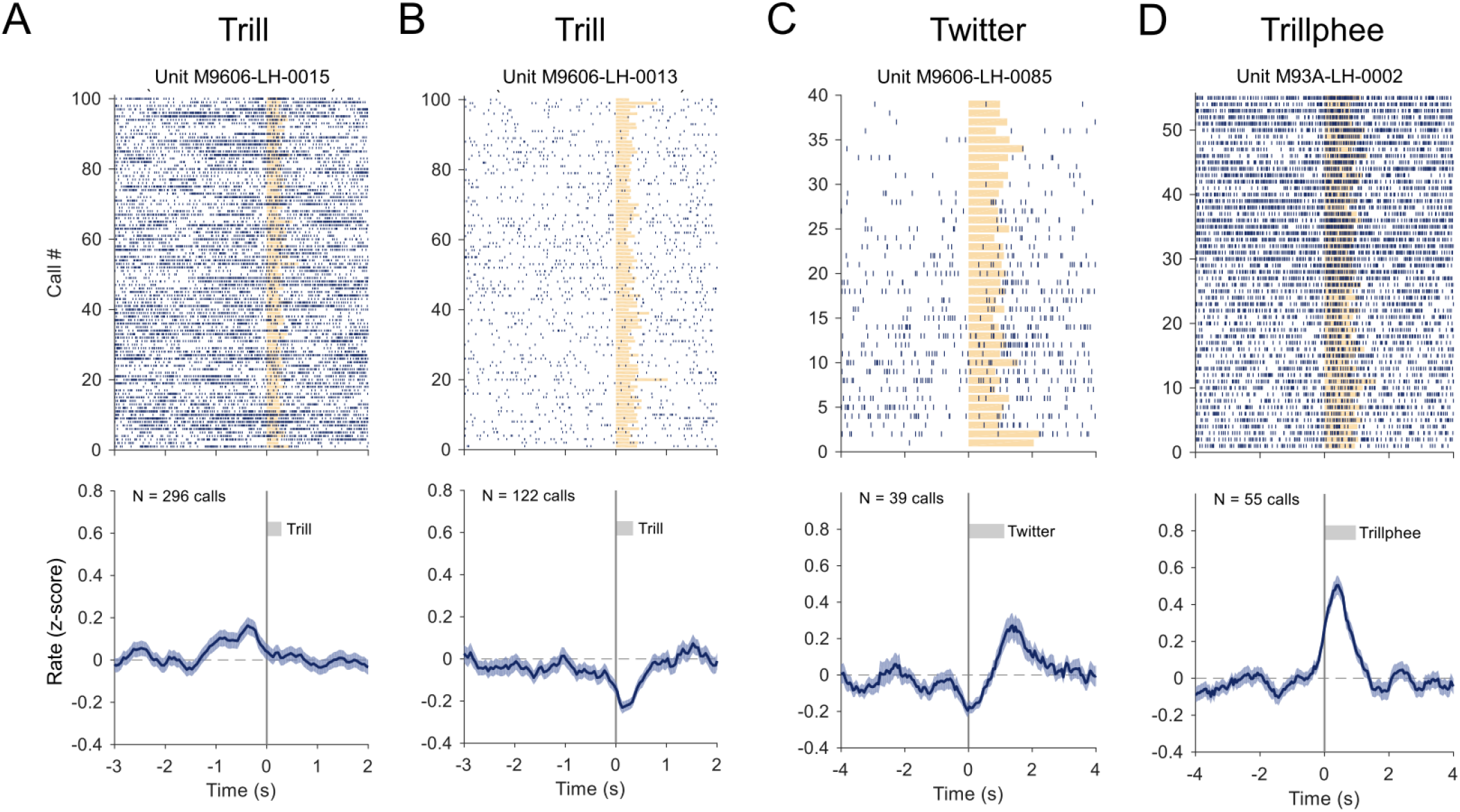
Example neurons showing modulation in activities for trill, twitter and trillphee calls. (**A**) Neuron showing increased activity before the vocal onset of trill calls. Top: Each vertical line indicates a spike. Spike timing is aligned to the vocal onset (time zero). Orange bars indicate the duration of trill calls. Only the first 100 calls are shown for clarity in the display (if the total number of calls is greater than 100). Bottom: Normalized firing rate (mean ± SEM). The firing rate is z-scored per trial and then averaged across trials (see Materials and Methods). N indicates the number of calls. The gray bar indicates the average duration of trill calls. (**B**) Neuron showing decreased activity during trill calls. (**C**) Neuron showing decreased activities near the vocal onset of twitter calls and increased activities after the end of twitter calls. (**D**) Neuron showing increased activities during the trillphee calls.

**Table 1.**
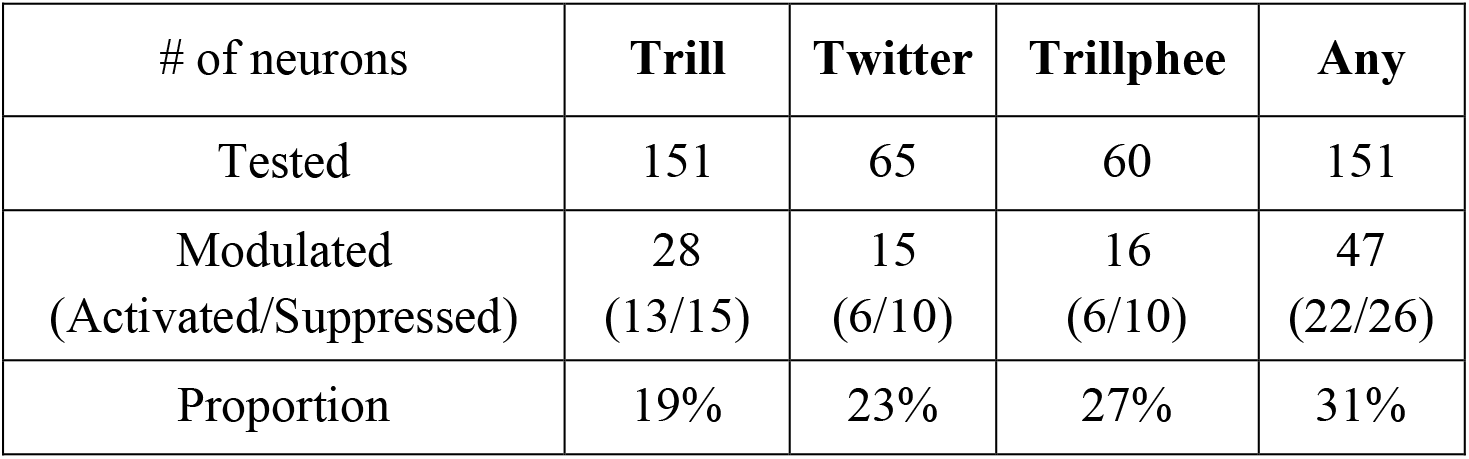
Number of neurons tested and modulated for each call type.

**Fig. 7.**
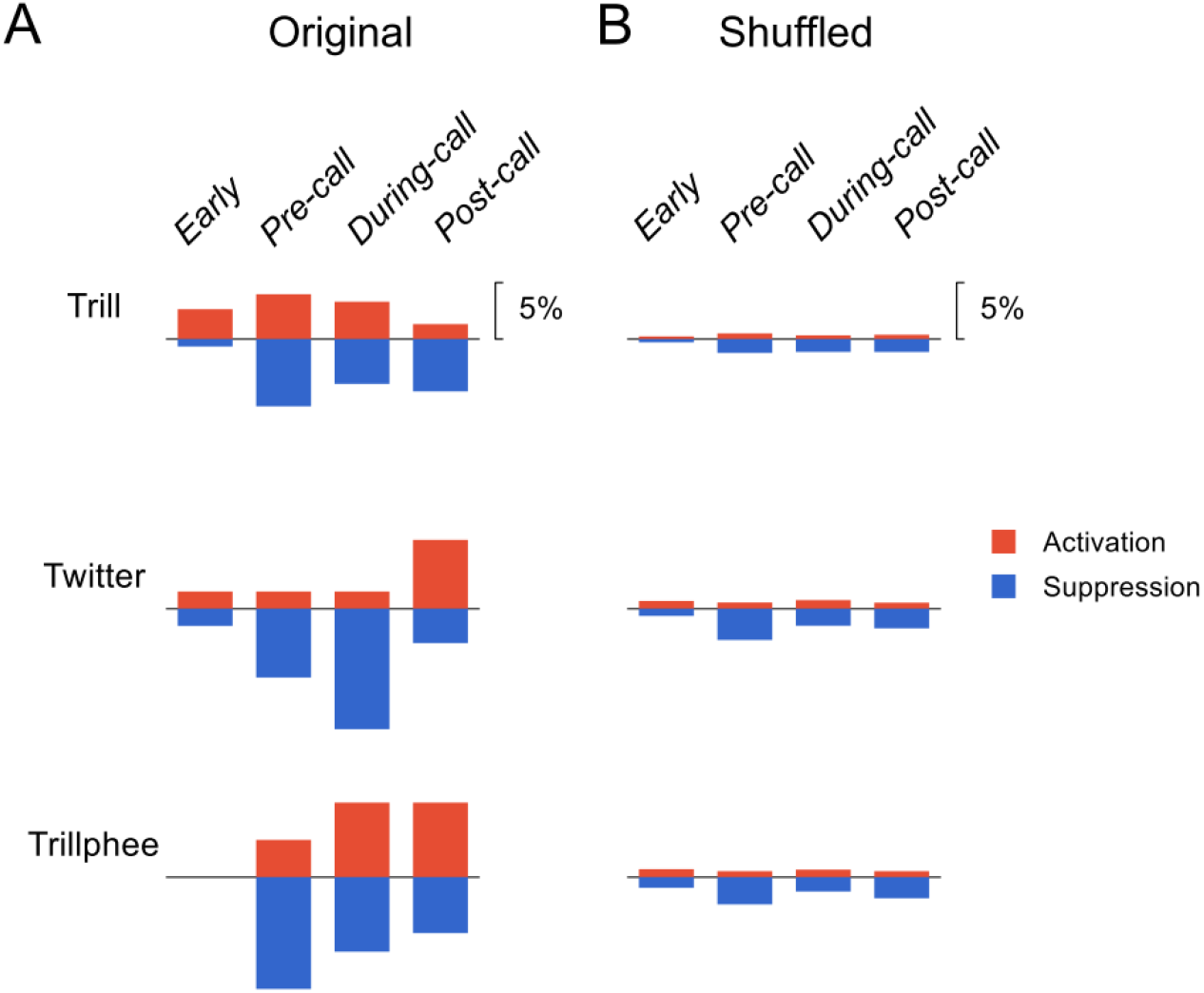
Proportion of neurons showing modulation in different analysis windows. (**A**) Comparison of the proportion of neurons showing activation (red) and suppression (blue) between four analysis windows and three call types: trill (top), twitter (middle) and trillphee (bottom). The analysis windows are labeled on top. The proportion is calculated with respect to the total number of neurons tested for each call type (see Table 1). (**B**) The same comparison as **A**, with shuffled data (see Materials and Methods).

### Distinct single neuron activities across call types

We then asked whether the activities in single neurons in the frontal cortex merely indicate the event of a vocalization or represent details of the motor production, i.e. distinct vocal structure in the different call types. The largest number of calls marmosets produced during social communication were trills. Twitters and trillphees were fewer. We therefore compared neural activities between trill and twitter calls and between trill and trillphee calls separately. Trills and twitters have very different acoustic structures. While trills are narrow-band calls with sinusoidal frequency modulations (Fig. 2B), twitters are wide-band calls with sharp upward frequency modulations in short syllables (Fig. 2C). If single neuron activities showed the same modulation for trills and twitters, then they were not likely to represent call type information. On the contrary, we found a subset of neurons showing different modulations between trills and twitters. One example neuron showed no modulation to trill calls but suppression during the production of twitter calls (Fig. 8A). Another example neuron showed activation for both trills and twitters (Fig. 8B). However, the activation for trills peaked before the vocal onset and the activation for twitters peaked near the end of calls. In total, 65 neurons were tested for both trills and twitters and 26 of them showed modulation for either type. As shown in Figure 8C, eleven neurons were modulated by trills but not twitters. Nine neurons were modulated by twitters but not trills. Six neurons were modulated by both but two of them showed different modulations. Therefore, 85% (22 out of 26) of modulated neurons showed a difference between the two call types. The fact that a majority of neurons showed a difference in the modulation between different call types suggested that the representation of call types in the marmoset frontal cortex existed in partially overlapping networks. Next, we compare the modulation for trills and trillphees. These two call types have similar acoustic structures, with the beginning part of trillphees resembles the structure of trills (Fig. 2D). The modulation for these two call types also showed a large difference. In total, 60 neurons were tested for both call types and 25 neurons showed modulation for either type (Fig. 8D). 88% (22 out of 25) of modulated neurons showed a difference between the two call types, including two neurons modulated by both types but with different modulations. These numbers indicate that even between call types with similar acoustic structures, there are distinct neuronal populations that represent each of them. Interestingly, nine neurons showed modulation only to trills but not trillphees, even if trillphees shared a similar acoustic structure with trills in the beginning part. This suggests that call type may be represented as unique categories rather than by its spectrotemporal features in a subset of neurons.

**Fig. 8.**
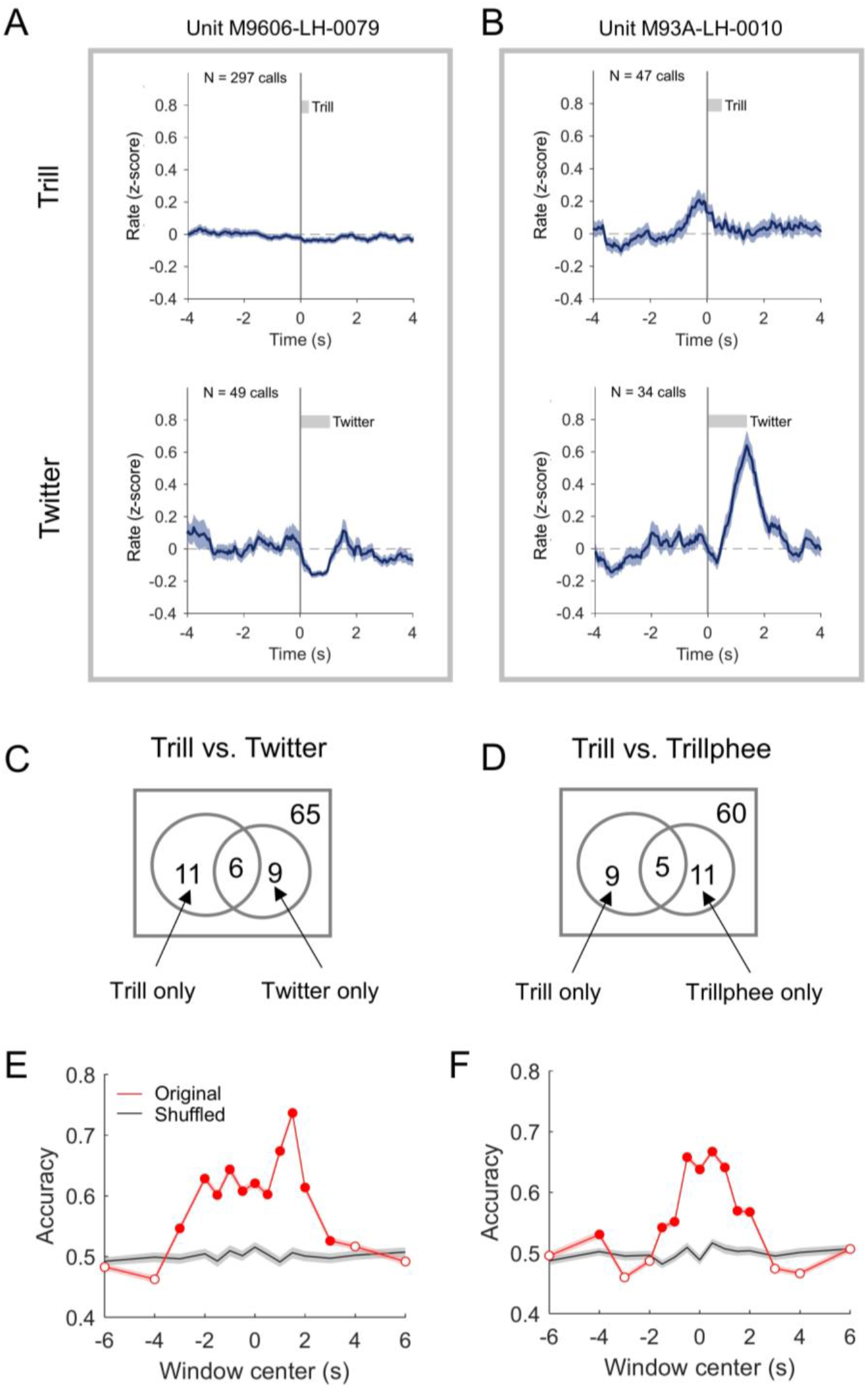
Distinct activity for call types in single neurons and in the neuronal populations. (**A**) Neuron showing different modulation between trill (top) and twitter (bottom) calls. The format of each panel is the same as **Fig. 6A** bottom panel. This example neuron showed no modulation for trill calls but decreased activities during twitter calls. (**B**) Neuron showing increased activities before the vocal onset of trill calls but near the end of twitter calls. (**C**) Venn diagram showing the relationship of neurons modulated by trills and neurons modulated by twitters. The number of neurons tested with both call types is indicated at the upper right corner of the rectangle. The circle on the left indicates the group of neurons modulated by trill calls. The circle on the right indicates the group of neurons modulated by twitter calls. The number in the overlapping region is for the neurons modulated by both call types. Two of these neurons showed differences in modulation between the two call types. (**D**) Same format as **C**, for neurons modulated by trills and neurons modulated by trillphees. Two of the five neurons with modulation to both call types showed differences in modulation between trill and trillphee. (**E**) Classification accuracy based on activities from neuronal populations to predict trill vs. twitter calls (see Materials and Methods). Red: original data; Black: shuffled data. Accuracy (mean ± SEM) is calculated using neural activities in a sliding window centered at different time points relative to vocal onset (time zero). Filled circle: accuracy significantly higher than that from shuffled data. Open circle: insignificant. (**F**) Same format as **E**, for predicting trill vs. trillphee calls.

Given the difference in modulation at the individual neuron level, we postulate that the call type for each vocal production can be predicted from the activities of the neuronal population. We used a linear classifier with Monte Carlo simulations to decode call types between trills and twitters for each trial, using the activities drawn from all neurons that were tested with these two call types (see Materials and Methods). The classifier was trained separately by data in a sliding window (one second in length) near vocal onset and the performance was evaluated by classification accuracy using testing data in the same window (Fig. 8E). We found that the accuracy was above chance level and was significantly increased from a null distribution obtained by shuffled data within about two seconds before and after vocal onset. Similar results were found when decoding call types between trills and trilphees (Fig. 8F). This suggests that population activities within a short period of time around a single vocalization can distinguish the type of call about to be produced or being produced. It is interesting that the classification accuracy significantly increased before vocal onset, suggesting preparatory activities from the frontal network.

## Discussion

We studied the neural mechanism of vocal production in the context of social communication for a highly vocal non-human primate species, the marmoset. In particular, we focused on a group of vocal signals used for marmoset communication, which have distinct acoustic structures and are often exchanged with conspecifics under different social contexts. We investigated the neural correlate of the production of these call types in the frontal cortex, a structure known to be important for human speech and communication. Overall, we provided three pieces of evidence for the functional role of the frontal cortex in generating different types of social communication calls in marmoset monkeys. First, we observed modulations in various regions in the frontal cortex in both LFP and single neuron activities when marmosets are generating different types of vocalizations involved in social communications with conspecifics. Second, we found differences in the pattern of neural activities for the major types of calls. This includes differences in modulations in LFP power, single neuron activities and population activities from neuronal groups. Third, theta-band LFP activities were found to have increased power during twitter calls and have phase lock to the individual syllables within twitters. These findings suggest an important function of the frontal cortex underlying the production of social communication signals in marmoset monkeys.

### Motor related beta oscillations in humans and monkeys

Oscillations in low-frequency neural signals have been known for a long time in both humans and non-human primates for motor movement. Beta-band power in the frontal cortex exists when the subject is at rest and it decreases when the subject starts to make voluntary movements (i.e. beta-band suppression), which occurred most prominently in sensorimotor cortex. This was demonstrated by a large number of previous studies, mostly involving hand movements and arm reaching tasks (35–37).

In our data, we observed widespread beta-band suppression for the production of all four types of calls across the recorded regions in the marmoset frontal cortex. Overall, the suppression started before the vocalization onset (for phee, trill and twitter calls) (Fig. 2, Fig. 3C). These observations are consistent with previous findings in other movement modalities (e.g. arm and hand movement), suggesting that the marmoset frontal cortex is modulated by the production of social communication calls. Furthermore, the magnitude and time course of beta-band suppression showed a difference across the four types of calls (Fig. 3). Trill call induced the smallest magnitude and shortest duration in suppression, whereas phee calls induced the longest duration. This matches closely with the vocal behavior in that trill is the softest and shortest call among the four call types and phee has the longest duration (31). Previous studies have found that beta-band suppression showed correlations with sensorimotor processing (39), motor preparation and initiation (40–42). Using grasping and reaching tasks, the pattern of beta-band modulation was found to be correlated with the specific types of movement (37, 43–45), duration (46), context (47, 48) and movement sequences (49). Our data on marmoset vocalization also showed correlations between the suppression and the motor output features, such as amplitude and duration, consistent with these findings. Interestingly, the start of beta-band suppression of phee calls occurred about half a second before vocal onset, much earlier than that of the other three call types (Fig. 3C). This may suggest that the production of phee calls in social communication involves extended preparation or cognitive process compared to the other three call types. Together, the difference in beta-band suppression across call types implies previously unknown cortical processing linked to the selection and production of vocal signals at a higher cognitive level during social communication.

### Theta-band activities in speech and vocal motor control

Rhythmic activities in the theta frequency band have drawn wide interests in human speech research. In most spoken languages, syllables occur with a repetition rate of 4-9 Hz (50), overlapping with the theta frequency range. Theta oscillations in the neural signals have been found in the human motor cortex and are thought to relate to sensorimotor processing in speech (51–53). Theta activation is also observed when subjects are making compensatory adjustments when auditory feedback is suddenly altered (54). In the motor control of hands, fine movements of fingers are found to be correlated with theta oscillations in the motor cortex (55). A recent animal study has found theta-band signals in the rodent motor areas associated with skilled movements that require coordinated motor sequence during reaching and grasping (56). In this rodent study, several key behavioral timepoints are found to be time-locked to the theta cycles (56). Here we observed theta-band activation in the frontal cortex of marmosets during vocal production. Surprisingly, the individual syllables of twitter calls are phase-locked to the theta oscillation cycles. To the best of our knowledge, this is the first time such observations are reported in non-human primate vocal behaviors.

The twitter calls of marmosets have an inter-syllable interval at about 140ms (7.1 syllables per second) (31). Other call types with repetitive structure, although less frequently produced, also have a syllable repetition rate around 4-10 Hz, such as peep-string (31) and fragmented phees (19, 21, 57). Empirically, the temporal structure of some marmoset call types resembles the syllable structure in human speech (4-9 Hz). Moreover, a recent study found that the mouth movement during marmoset vocal production is following the rhythm of the acoustically defined syllables (57). The authors proposed that coordinated motor control for articulatory (e.g. mouth) and phonatory (larynx) systems give rise to the rhythmic vocal output in marmosets (57). Such coordination is found to be crucial for human speech production. The theta oscillation found here, which is tightly coupled to the instantaneous vocal output, is likely to underly the control for the generation of syllables in marmoset calls and may also provide a neural basis for the coordination of articulators. Interestingly, a large portion of previous research on periodic movement patterns attributes the neural control to the “central pattern generator” in the brainstem. In contrast, we found theta-band oscillation in the frontal cortex, a region likely to be associated with voluntary motor control, suggesting the possibility of a high-level control mechanism in marmoset vocalizations. It remains an open question whether there are similar oscillation signals in the brainstem and where the cortical oscillation is originated. The fact that marmoset calls and human speech have similar syllable rates and the theta activities found in the frontal cortex suggests that there may be a common precursor in the evolution of monkey vocalization and human speech for the neural basis of temporal structures.

### Comparison of single neuron results to previous studies

Comparing our results here to our previous experiment in the behavioral chamber (28) and others’ data (29) on phee call production, we noticed some differences in the modulations in the neurons in the frontal cortex. For example, there seems to be a fewer proportion of neurons modulated in vocal production in the social environment. While 40%-50% of neurons were found to be modulated in the two previous studies in an isolation environment, only about 30% were modulated in this data. Also, it seems that the size of modulation in the modulated neurons are smaller in this data for trill/twitter/trillphee compared to that for phee calls tested in the behavior chamber, although this is hard to be precisely quantified given the different method used in normalizing neural activities and settings of analysis windows. Here we do not know how the same neurons in the colony experiment would be modulated by phee calls in the behavior chamber, but the difference in modulation across call types reveals potentially different mechanisms underlying the production for different call types in the marmoset frontal cortex, as previous studies have suggested that premotor cortex likely represents movement patterns (58–60). It may also be caused by different contexts which give rise to a global modulation to neuron’s firing rate (61).

The time period of modulation in our observation for trill, twitter and trillphees is more sophisticated than what we saw for phee calls. For phee calls in the isolation environment, most modulations occurred in a pre-vocal period about half a second before vocal onset or during vocalization (28, 29). Some neurons showed activation right after vocal production as well (see Fig. 2 in (28)). Interestingly, in our current experiment, we saw modulation much earlier than the previous pre-vocal period (corresponding to the pre-call window here, see Materials and Methods). Some neurons showed activations within the [−2, −0.5]s window for trill calls (Fig. 6A). For both trill and twitter, we also saw activation near the end of calls. For trillphee, the strongest activation only occurred during vocal production. For all of trill, twitter and trillphee, suppression almost always overlapped with the entire vocal production. It will be an interesting question for future studies regarding the functional role of modulations at different time windows.

### Influence of the free moving paradigm and the social communication contexts

Another factor that will largely affect the analysis of modulation is baseline activities. In traditional motor experiments, animals were usually restrained in a primate chair and instructed by visual cues. Before the cue and movement, one could assume there is no or little movement of the animal or cognitive processes for potential behavior. Unlike these paradigms, experiments with freely moving animals do not have an absolute baseline period in which one can assume no motor-related activities. In this case, we would expect a lot more variations in the firing rate in the baseline window. The analysis here relies on the temporal alignment of particular behaviors as the trial onset. In vocal production, in order to see possible modulation of vocal events, we aligned the neural activities to the onset of vocalizations and assumed that other activities are not time-locked (or with a fixed delay) with vocal production. Essentially, we expect that modulation induced by non-vocal behaviors gets averaged out in our baseline windows. Of course, this method has limitations. If the variations in neural activities induced by other behaviors are larger in the baseline window than the modulations induced by vocalizations in analysis windows, then it would be difficult for the vocal-induced modulations to be identified, or it requires a substantially large number of trials to average out the “noise” from other behaviors. When comparing the behavior of marmosets in the colony with the one in the behavioral chamber, we noticed that there are usually moderate locomotion activities when animals were not vocalizing in the chamber, whereas there are a lot more activities in the colony, including locomotion, food and water intake, grooming, etc. Therefore, it is not surprising that the variation of neural activities is overall larger than that in a behavioral chamber. Given this fact, we think the number of vocal modulated neurons in the colony experiment is likely to be underestimated.

### Implications to vocal control in non-human primates

Vocal communication through speech is probably the most intriguing behavior that distinguishes human intelligence from animals. It has been a long-standing question of how the brain controls the generation of speech and other communication signals (62). The general notion has been that the lateral part of the cortex, including frontal, parietal and temporal regions are involved in speech production and learning, whereas the medial structures in the cortex and the brainstem, including ACC and PAG, are involved in emotional vocalizations, such as cry and laughter (4). Studies in non-human primates, however, have generated controversial results in the past decades. Experiments using electrical stimulation and conditioned vocalizations seemed to suggest that the lateral frontal cortex of monkeys was dispensable for vocal production (11, 63, 64). Recent experiments with single neuron recordings in macaques with operant conditioning paradigm found activities in the premotor and prefrontal cortex for conditioned vocalizations but not for spontaneous vocalizations (13, 14). Studies using freely moving marmoset monkeys, on the other hand, have observed neural activities in the frontal cortex during self-generated phee calls (28, 29). These studies provided initial evidence for the involvement of the lateral frontal cortex in vocal production.

Modulation of neural activities by different call types has not been studied in the frontal cortex before. Previous studies on the brainstem have found separate populations of neurons that are correlated with the production of several call types in squirrel monkeys (65–67). In squirrel monkey PAG, a subset of neurons was found to be only activated to one or a subset of vocal types (67). In ventrolateral pontine areas (VOC), which receive input from PAG and project to several cranial motor neuron pools involved in phonation, neurons were found to be only modulated by frequency-modulated (FM) calls but not non-FM calls (65). In the downstream motor neuron pools, more than half of the neurons showed activities only to frequency modulated call types while another subset showed activities to both FM and non-FM calls (66). These activities were thought to be related to the “central pattern generator” (68). Further, anatomical tracing studies for PAG seemed to suggest that several upstream regions, including the hypothalamus and anterior cingulate cortex, may play a role in driving the different neuronal populations in PAG that were activated for different call types (69). However, pharmacological inactivation in PAG was found to abolish vocal fold activity induced by the cingulate cortex but not by the laryngeal motor cortex (70), which provided evidence for two separate pathways for vocal fold control, i.e. one from the limbic cortex and the other from the neocortex. A recent study using retrograde tracing in marmoset laryngeal muscles revealed frontal cortical projections from the premotor and primary motor cortices (71). In our experiment, we tested LFP and individual neurons’ modulations for different call types in the frontal cortex. The fact that both LFP and individual neurons showed distinct activities for different call types suggests that these activities do not simply act as an initiation command or timing signal for brainstem structures, but rather contribute to the generation of the acoustic structure in the individual vocal signals. Therefore, our study provides support for a crucial function of the pathway involving the lateral frontal cortex, i.e., the category and feature of the vocal signals may be shaped by activities from the premotor or primary motor cortices, rather than being dictated by limbic or brainstem activities.

Interestingly, most neurons showing modulations in vocal production were modulated only by one out of two call types. A subset of neurons was modulated only by trill calls but not by trillphee calls (Fig. 8D), despite the similarity in the frequency modulation in the call structure. A further hypothesis can be proposed that there exists an abstract representation of call type categories in a subset of neurons in the lateral frontal cortex. Future studies are needed to investigate how these neurons communicate with other neurons in the frontal cortex and in the brainstem nuclei.

In summary, our study provides new evidence for the role of the frontal cortex in the vocal production of social communication calls in marmosets. It implies that marmosets may engage sophisticated vocal behaviors during communication through neural mechanisms based in the lateral frontal cortex.

## Materials and Methods

For details regarding the rationale of design, vocal behavior, acoustic and neural recording techniques, and histology, please refer to the Supplementary Methods (SI Appendix).

### Animal preparation and experimental procedure

Two adult marmosets were used in the experiment (M9606, M93A, both are male). M9606 was implanted with a 16-channel electrode array (Warp-16, Neuralynx) and M93A was implanted with a 32-channel electrode array (Warp-32, Neuralynx). Both arrays were in the left hemispheres. Implantation of the arrays followed a two-step procedure established in the lab previously. First, marmosets underwent surgical procedures and were implanted with a head cap (72). After the marmosets were fully recovered from the surgery, a craniotomy was created in the frontal areas exposing part of the premotor and primary motor cortices. The location of the craniotomy was identified with reference to the marmoset brain atlas (73), using the lateral sulcus as a surface landmark. The array was positioned above the dura and sealed by Silastic (Qwik-Sil, WPI) and fixed by dental acrylic. Tungsten electrodes (4-12MΩ, FHC or A-M Systems) housed in the arrays were advanced through the dura and adjusted in small steps (~50μm) to search for single neurons. All experimental procedures were approved by the Johns Hopkins University Animal Care and Use Committee and were in compliance with the NIH guidelines.

At the beginning of the experiment each day, the experimental subject was brought to a custom-built RF/EMI shielded chamber (74) and head-fixed in a primate chair. Neural signals were checked using the wireless recording system. A polycarbonate protection cap was then mounted on the marmoset’s head to protect the wireless headstage. The marmoset was then moved into the colony. The subject M9606 was recorded in a shielded booth located at the corner of the colony room. An analog wireless system (see Supplementary Methods for details, SI Appendix) was used for early sessions and a digital wireless system was used for later sessions (LFP signals were only obtained from these later sessions). A parabolic-reference microphone pair was placed 50 inches in front of the plastic cage (see Supplementary Methods for details, SI Appendix). The subject M93A was recorded in its home cage. Before recording, the cage was moved to the side of the colony room next to the wall. Acoustic foams were placed in front of the wall near the cage to reduce sound reflections. The digital wireless system was always used for this subject. The parabolic-reference microphone pair was placed 40 inches in front of the home cage. To synchronize the neural and acoustic recordings, a pulse train with a period of two seconds was sent to the neural and acoustic recording devices simultaneously and was used to align the timing of the two sets of signals. A typical session lasted 2-5 hours long. After recording, the marmoset was moved back to the chamber. The wireless headstage was removed, electrodes were advanced and the marmoset was sent back to the colony.

### Data analysis

Pre-processing was first performed including segmentation of vocalizations in the acoustic recordings, classification of call types, spike sorting, and artifact detection (see Supplementary Methods, SI Appendix). Calls overlapped with artifacts were excluded from subsequent analysis. To ensure temporal alignment of neural signals, any calls with a duration shorter than 200 ms were also excluded. This was done to prevent neural modulations from being washed out by averaging weak and transient activities for short trill calls together with more pronounced activities for longer calls.

The time-frequency representation of LFP power was calculated using the FieldTrip toolbox (75). The spectrogram of LFP signals in each trial from a single electrode channel was first calculated by a wavelet transformation (Morlet wavelets with a width of 7) and then averaged across all trials (including all calls available for that channel) (76, 77). We chose a baseline window of [−3, −1] sec relative to the vocal onset to normalize LFP power. In the time-frequency representation, the power at each time point is normalized to the average power within this baseline window for each frequency band respectively. To calculate LFP power in a particular frequency band, the raw LFP signals were first band-pass filtered (12-30 Hz for beta band; 4-8 Hz for theta band) and then converted to analytical amplitude by a Hilbert transform. The signals were further down-sampled to 200 Hz and the power was normalized with respect to the baseline window. To find the earliest time at which LFP power showed significant suppression or activation, we used a sliding window of 100 ms long and compared the LFP power in this window to that in the baseline window. We selected the mid-point of the earliest window in which LFP power showed significant difference (signed-rank test) and was more than two standard deviations away from the power in the baseline window (see below for the details of the standard deviation calculation).

To characterize any modulation in LFP power in beta- or theta-band for a given electrode, an analysis window of [−0.1,0.2] sec relative to the vocal onset was used to quantify the LFP power. A recording site (or channel) was significantly modulated by a call type if the LFP power in the analysis window was significantly different from that in the baseline window (signed-rank test) and was more than two standard deviations away from the baseline LFP power.

It is worth noting that using the standard deviation in the baseline LFP power as a criterion for modulation may induce bias, since the number of calls for each call type was largely different (e.g. there are a lot more trill calls than the three other call types), which would affect the size of standard deviation. To overcome this issue, we randomly drew 200 trials for each call type to calculate the averaged LFP power in the baseline window. We then used 200 ms non-overlapping windows (length comparable to that of the analysis window) to segment the averaged LFP power and calculated the standard deviation of the LFP power in these segments. This procedure was repeated 1000 times to bootstrap the mean standard deviation which was used as the criterion mentioned above.

To quantify the phase lock of twitter syllables to theta-band LFP, we calculated vector strength (78). The higher the vector strength, the stronger the phase lock was shown. Rayleigh statistics was used to test whether the phase angle in the theta oscillation at which each twitter syllable started has a non-uniform distribution.

For single neuron analysis, we included neurons with at least 15 calls for subsequent analysis. To reduce the variation of baseline firing rate across trials, we z-scored the firing rate for each trial (relative to a 20-sec window centered at vocal onset). Firing rates were calculated in 50 ms time bins. To characterize the modulation of spike activities, we used four analysis windows (time relative to vocal onset): [−2, −0.5] sec (early activities); [−0.5, 0] sec (pre-call activities); [0, 80% of call duration] (during-call activities); [80% of call duration, 0.5 sec after call end] (post-call activities). Firing rate in each of these windows was compared to that in a baseline window ([−8, −4] sec). A neuron is showing modulation in a specific analysis window if the firing rate in the analysis window is significantly different from that in the baseline window (signed-rank test) and is at least two standard deviations away from the baseline firing rate. Similar to LFP analysis, we used a bootstrap strategy to estimate the standard deviation of the baseline firing rate (using 500 ms segments). To get the number of modulated neurons with shuffled data, we circular-shifted the spike timing for each trial with a random amount and then calculated the mean firing rate across trials. The number of neurons with significant modulation with this shuffled data was calculated. This procedure was repeated 100 times to obtain the mean number of neurons with significant modulations (used in Fig. 7B).

For the population analysis, since the number of neurons recorded simultaneously during each session was small and the number of calls collected for each call type was different, we used a Monte Carlo simulation to balance the sample size. For a given neuron, twitter and trillphee calls may not have enough samples to compare with each other. Therefore, we built two types of classifiers to decode trill vs. twitter and trill vs. trillphee calls, respectively. For each type of classifiers, we included neurons with at least 20 calls to each call type and use the mean firing rate within a one-second long window centered at different time points relative to vocal onset to run the analysis (the classifiers are trained separately at each window). We randomly select 75% of trials from each call type to form the distribution of training data and the rest 25% to form the distribution of testing data. We then redraw these trials 5000 times for each call type to construct the actual training and test data set. A linear discriminant analysis method was applied to classify the two call types. This procedure was repeated 100 times to obtain the mean and standard error of the classification accuracy. For the shuffled data, we performed the same procedure except that we shuffled the call type labels during training.

Signed-rank tests were used to test the significance of neural modulations. One-way ANOVA, Kruskal-Wallis tests and *post hoc* analysis with the Bonferroni correction were used to compare the modulation size or time across multiple call types. A two-sample t-test was used to test the significance of the decoder accuracy. Significance was determined at an α-level of 0.05.

## Acknowledgment

We thank Reza Shadmehr and Cynthia Moss for helpful discussion on the data; Nathaniel Sotuyo, Shanequa Smith, Alexandra Prado and Jessica Lynch for assistance with animal care; Kristina Nelsen, Marcello Rosa and Brian Lee for help with histology and anatomical reconstruction; Calvin Qian, Emile-Victor Kuyl, Kevin Zhu, Hu Yi, for assistance with experiments and data preparation; Haowen Xu for optimizing the algorithm for call detection and classification. This work is supported by the National Institutes of Health (Grant DC 005808).

## Supplementary Methods

### Study design

Several components need to exist in order to study marmosets’ natural vocal communication in a social context: (a) marmosets being recorded in a freely-moving behavioral state to allow natural vocal production; (b) sufficient social contexts to elicit different types of social calls without conditioning or experimental reinforcement; (c) techniques to record the neural activities and the vocal signals continuously and reliably from the marmoset without interfering its behavior. These components require a systematic design in the experiment combined with a series of new techniques. Our lab has previously developed a behavioral paradigm to study natural vocal exchanges in a controlled experimental chamber (26), designed apparatus to allow wireless single-unit recordings in freely-moving marmosets (34), and utilized tethered recording system in the marmoset colony to study the auditory cortex during vocal production and perception (79).

In this experiment, we developed new techniques based on the existing ones to achieve the goal of our study. We performed our recordings in the marmoset colony, which provided an enriched social context. During the period of this study, the colony room housed about 40-50 marmosets including breeding pairs and families with adults and babies. The recordings from the experimental subject were done either in its home cage or in a specially designed shielded booth. In either case, the subject was able to see other marmosets in the room and participate in vocal exchanges with them. While this environment provided the social contexts to elicit all types of vocalizations from the marmoset, it poses challenges to both acoustic and neural recordings. We will detail the solutions in the following paragraphs.

### Vocal communication behavior

Previous studies have shown that marmosets utilize more than ten types of calls in social communication both in the field (32, 33) and in the lab setting (31). Four of these types (phee, trill, twitter, and trillphee) are most frequently produced and have been well quantified in terms of their acoustic structures (31, 80). In the marmoset colony, individual animals housed there constantly exchange vocalizations in and outside of their home cages. The experimental subject, either tested in our recording apparatus (a shielded booth, as detailed below) or in its home cage, maintained vocal interactions with other individuals in the room. They produced the same range of call types as they are not being recorded and as in the census of general vocal repertoire in the marmoset population. The experimental subject either spontaneously generated these calls or in exchange with other individuals housed in the same room.

### Targeted acoustic recordings

Unlike a dedicated recording chamber, the marmoset colony does not have the sound isolation power ideal for acoustic recordings. Sound targets (individual marmosets) are relatively close to each other in spatial locations. Vocalizations produced span a wide dynamic range and are often overlapping in time. Figure 1F (middle) illustrates an example recording from the colony, where vocalizations from multiple marmosets and background noise are captured by the same microphone. The goal of the targeted acoustic recording is to overcome this challenge and obtain clean recordings of the vocalizations from the experimental subject (target).

The criteria for clean recordings should include the following. First, both loud and weak calls can be recorded. Since marmoset vocalizations have a relatively wide dynamic range, with some loud phees over 100dB SPL and some trills below 50dB SPL (81), the recording system needs to ensure that the weak calls will not be buried in background sounds and become un-detectable. The background sounds include two categories. One is room noise, from the air circulation system, marmosets’ non-vocal activities in cages (e.g. jumping), and occasional human activities. The other is vocalizations from other marmosets. For example, a trill call from the target marmoset can easily be masked by a loud phee call from a neighboring marmoset. Second, the recording system should enable clear separation of vocalizations between target marmosets and non-target marmosets (source separation). For example, if a target marmoset is silent and a non-target neighbor makes a loud call, this call should not be mixed into the target marmoset’s vocalizations. Third, the recording system should facilitate precise and efficient segmentation during post-processing. Because of the large number of vocalizations generated and recorded in the colony, there is usually a heavy burden in the post-processing to segment calls and classify call types. It is favorable if the hardware design can reduce the load of post-processing. The key to fit these criteria is to increase the signal-to-noise ratio of the target marmoset vocalizations.

Traditional techniques in acoustic recordings have limitations in this scenario. For example, directional microphones may reject sound coming from the side but may not provide enough differentiation for sources in the front, especially when individual marmosets are housed in cages next to each other. One can presumably move the experimental subject to a separate cage far from its neighbors. However, this is less desired since the social context is changed as the distance increases between the experimental subject and other individuals in the room. Collar devices, small microphones mounted on an attachment to the animal, may obtain decent sound intensity from the experimental subject, but they require adaptation for the animal and more manual labor to attach them and to change the battery, causing interference to the recording sessions and potentially the subject’s behavior.

Here we designed a parabolic-reference microphone pair to solve the recording problem (Fig. 1D). A parabolic microphone is composed of a parabolic reflector and a microphone placed at the focal point, which is often used in field recordings (82). It selectively amplifies signals coming from the front and has a higher gain in high frequencies than in low frequencies. A reference microphone is the same microphone model used in a traditional way without a reflector, placed next to the parabolic microphone (both directed at the sound target). We used a parabolic reflector with a diameter of 20.5 inches, a depth of 6 inches and a focal length of 4 inches, made of transparent polycarbonate (generic supplier from eBay). A microphone (AKG C1000S, cardioid pick-up pattern) was pointed towards the reflector with its diaphragm placed at the focal point of the reflector. Theoretical calculations of the acoustic properties of the parabolic reflector (83) revealed that the lowest frequency cutoff for recording is about 650Hz. Since the fundamental frequency of most marmoset vocalizations is above 5kHz (31), this parabolic microphone is well suited for the frequency range of marmoset vocalizations.

The pick-up pattern (the relationship between gain and angle) of the parabolic microphone is tested in a sound-attenuating chamber. The gain of the parabolic microphone was measured with pure tones of different frequencies (1kHz, 2kHz, 4kHz, 8kHz, 16kHz) played from a speaker (KEF LS50) at different angles (0°-180°, with 30° step; 0° incident angle means facing directly to the speaker) and was compared to the reference microphone (Fig. 1E, data for 8kHz, interpolated by a spline function). The parabolic microphone (Fig. 1E, green curve) has a much sharper pick-up pattern than the reference microphone does (Fig. 1E, brown curve). For sounds coming from the front, the parabolic mic has nearly 20dB additional gain than the reference microphone. As the sound source offsets in direction, the gain of the parabolic microphone quickly drops and then becomes smaller than the reference microphone. While a traditional microphone (used as the reference microphone) achieves about 10dB front and back gain difference, the parabolic microphone achieves over 30dB of such difference. This feature suggests two things. One is the parabolic microphone has superior directionality to a traditional microphone; the other is by comparing the intensity of signals recorded by the parabolic microphone and reference microphone, one can identify whether the source is located in the target direction.

In our recording setup, we placed this microphone pair in front of a target marmoset cage and calculated the difference of signal intensity between the two microphone channels. Given the dimension of the cage, the distance from the cage to the microphone and the pick-up pattern, we obtained the threshold of intensity difference. If the signal intensity in the parabolic channel is higher than that in the reference channel by an amount larger than the threshold, the source is from the target marmoset cage. Figure 1F (top and middle) shows an example recording clip from the two channels. While both microphones pick up vocalizations and noises in the room, signals from the reference microphone resemble a uniform background, whereas signals from the parabolic microphone enhance target vocalizations, as they are from the direction the microphones were faced at. Room noise and vocalizations from non-target locations have similar intensities in the two microphone channels, as they are mostly coming from the side. When subtracting the two spectrograms (Fig. 1F, bottom), room noise and vocalizations from non-target marmosets canceled out and only the vocalizations from the target marmoset were left (Fig. 1F, bottom, red circles). Using this setup, we can enable reliable detection of target vocalizations even when they are weak in intensity or overlapped by calls from other marmosets.

After recording, a custom-written Matlab program was used to segment target vocalizations from the continuous recordings based on machine learning algorithms. Call types were classified by the same program. All time points of vocalizations and call type labels were checked and corrected by human experimenters by visual inspection of the spectrograms.

### Wireless neural recording in the colony environment

Wireless neural recording techniques have previously been applied in the lab in a radio frequency (RF) shielded behavioral chamber (34). We adapted this technique in the colony recording in a two-step process. In the first step, we used an analog wireless system (W16, Triangle Biosystems, or TBSI) in a custom-built RF-shielded booth to provide a supporting environment for reliable RF transmission (Fig. 1A). The booth is built with copper mesh and lined with RF absorption foam (EHP-5CV, ETS Lindgren) on most of the sidewalls and ceilings. The lower front part of the cage has no foam installed so the subject can see the colony through the booth. The experimental subject is placed in a plastic cage (60cm x 41cm x 30cm) inside the booth made of plexiglass and nylon mesh. The wireless receiver is fixed directly above the plastic cage. The plastic cage is transparent to wireless signals and the shielded booth isolates the environment from interference in the colony room and reduces reflection of RF transmission from the walls of the booth. We tested this setup using the wireless headstage and a spectrum analyzer (R3172, Advantest). Within the space of recording, we obtained clean, reliable wireless transmission with stable signal strength, regardless of the headstage location or orientation, indicating there is no interference from outside of the booth or from multi-path propagations inside the booth.

In the second step, we used a digital wireless system (W2100-HS32, Multichannel Systems, or MCS) to allow for recordings when the subject is free-roaming in its home cage (made with metal mesh and panels) (Fig. 1B). Digital transmission is known to be more robust to noise interference than analog transmission. Therefore, the recording location does not have to be protected by RF shielding. In our experiment, the receiver was placed above the subject’s home cage. Before a recording session started, the orientation of the antennae was adjusted so that the real-time transmission quality was at an excellent level (indicated in the Multi Channel Experimenter Software, MCS). This transmission quality was monitored throughout the recording sessions. In the rare case when there was data loss during transmission, the timestamps were logged by the recording software and the neural activities near that time period were excluded from the analysis. We used a custom-built battery pack (600mAh, ~11g) mounted on the headstage, which supported a continuous recording of 32 channels for up to 5 hours.

For the analog system, raw neural signals were amplified (Lynx-8, Neuralynx), band-pass filtered (300-6000Hz) and digitized at a 20kHz sampling rate (PCI-6071E, National Instrument). Data were stored on a recording computer through a custom-written Matlab program. For the digital system, raw neural signals were band-pass filtered for spikes (300-6000Hz) and the local field potential (LFP, 1-300Hz) respectively and stored on a recording computer through a software of the wireless system (MC_Rack or Multi Channel Experimenter, MCS) at a sampling rate of 20kHz. LFP signals were then down-sampled to 1kHz for analysis. Spike waveforms were sorted off-line with a template matching method in a custom-written Matlab program (28, 34). Neurons recorded from the same electrodes on the same day were considered the same units. Any artifact in spike or LFP signals was detected automatically in preprocessing and excluded from the analysis.

### Histology

When all experiments were finished, electrolytic lesions were made by passing a small DC current through some of the recording electrodes (25μA, 25s). The animals were anesthetized by ketamine and euthanized by pentobarbital sodium. It was then perfused with a phosphate-buffered solution and 4% paraformaldehyde. Subject M9606’s brain was sectioned in the coronal plane. A series of staining methods were applied to distinguish cortical regions, including Nissl, Myelin and SMI-32 immunohistochemistry stains. To identify the locations of recording electrodes, the stained sections were scanned into digital images and reconstructed into a 3D model by comparing them to the standard marmoset brain atlas. The cortical regions were then segmented with reference to the standard atlas. The lesion marks were visually identified and assigned to one of the cortical regions in the segmented model.

## References

1. Cheney DL, Seyfarth RM (2018) Flexible usage and social function in primate vocalizations. Proc Natl Acad Sci U S A:201717572.

2. Roy S, Miller CT, Gottsch D, Wang X (2011) Vocal control by the common marmoset in the presence of interfering noise. J Exp Biol 214(Pt 21):3619–29.

3. Takahashi DY, Narayanan DZ, Ghazanfar A a (2013) Coupled oscillator dynamics of vocal turn-taking in monkeys. Curr Biol 23(21):2162–8.

4. Pisanski K, Cartei V, McGettigan C, Raine J, Reby D (2016) Voice Modulation: A Window into the Origins of Human Vocal Control? Trends Cogn Sci 20(4):304–318.

5. Picardo MA, et al. (2016) Population-Level Representation of a Temporal Sequence Underlying Song Production in the Zebra Finch. Neuron 90(4):866–876.

6. Flinker A, et al. (2015) Redefining the role of Broca’s area in speech. Proc Natl Acad Sci 112(9):2871–2875.

7. Bouchard KE, Mesgarani N, Johnson K, Chang EF (2013) Functional organization of human sensorimotor cortex for speech articulation. Nature 495(7441):327–32.

8. Conant DF, Bouchard KE, Leonard MK, Chang EF (2018) Human Sensorimotor Cortex Control of Directly Measured Vocal Tract Movements during Vowel Production. J Neurosci 38(12):2955–2966.

9. Chartier J, Anumanchipalli GK, Johnson K, Chang EF (2018) Encoding of Articulatory Kinematic Trajectories in Human Speech Sensorimotor Cortex. Neuron 98(5):1042–1054.e4.

10. Dichter BK, Breshears JD, Leonard MK, Chang EF (2018) The Control of Vocal Pitch in Human Laryngeal Motor Cortex. Cell 174(1):21–31.e9.

11. Jürgens U (2002) Neural pathways underlying vocal control. Neurosci Biobehav Rev 26(2):235–258.

12. Jürgens U (2009) The neural control of vocalization in mammals: a review. J Voice 23(1):1–10.

13. Coude G, et al. (2011) Neurons controlling voluntary vocalization in the macaque ventral premotor cortex. PLoS One 6(11):e26822.

14. Hage SR, Nieder A (2013) Single neurons in monkey prefrontal cortex encode volitional initiation of vocalizations. Nat Commun 4:2409.

15. Eliades SJ, Miller CT (2017) Marmoset vocal communication: Behavior and neurobiology. Dev Neurobiol 77(3):286–299.

16. Takahashi DY, et al. (2015) The developmental dynamics of marmoset monkey vocal production. Science 349(6249):734–8.

17. Gultekin YB, Hage SR (2017) Limiting parental feedback disrupts vocal development in marmoset monkeys. Nat Commun 8:14046.

18. Takahashi DY, Liao DA, Ghazanfar AA (2017) Vocal Learning via Social Reinforcement by Infant Marmoset Monkeys. Curr Biol 27(12):1844–1852.e6.

19. Pomberger T, Risueno-Segovia C, Löschner J, Hage SR (2018) Precise motor control enables rapid flexibility in vocal behavior of marmoset monkeys. Curr Biol 28(5):788–794.e3.

20. Zhao L, Rad BB, Wang X (2019) Long-lasting vocal plasticity in adult marmoset monkeys. Proc R Soc B Biol Sci 286(1905):20190817.

21. Zhao L, Roy S, Wang X (2019) Rapid modulations of the vocal structure in marmoset monkeys. Hear Res 384. doi:10.1016/j.heares.2019.107811.

22. Chow CP, Mitchell JF, Miller CT (2015) Vocal turn-taking in a non-human primate is learned during ontogeny. Proc R Soc London B Biol Sci 282(1807).

23. Snowdon CT, Elowson a. M (1999) Pygmy Marmosets Modify Call Structure When Paired. Ethology 105(10):893–908.

24. Rukstalis M, Fite JE, French JA (2003) Social change affects vocal structure in a callitrichid primate (Callithrix kuhlii). Ethology 109(4):327–340.

25. Elowson AM, Snowdon CT (1994) Pygmy marmosets, Cebuella pygmaea, modify vocal structure in response to changed social environment. Anim Behav 47(6):1267–1277.

26. Miller CT, Wang X (2006) Sensory-motor interactions modulate a primate vocal behavior: antiphonal calling in common marmosets. J Comp Physiol A Neuroethol Sens Neural Behav Physiol 192(1):27–38.

27. Miller CT, Beck K, Meade B, Wang X (2009) Antiphonal call timing in marmosets is behaviorally significant: interactive playback experiments. J Comp Physiol A Neuroethol Sens Neural Behav Physiol 195(8):783–9.

28. Roy S, Zhao L, Wang X (2016) Distinct neural activities in premotor cortex during natural vocal behaviors in a New World primate, the common marmoset (Callithrix jacchus). J Neurosci 36(48):12168–12179.

29. Miller CT, Thomas AW, Nummela SU, de la Mothe LA (2015) Responses of primate frontal cortex neurons during natural vocal communication. J Neurophysiol 114(2):1158–71.

30. Nummela SU, Jovanovic V, de la Mothe L, Miller CT (2017) Social context-dependent activity in marmoset frontal cortex populations during natural conversations. J Neurosci 37(29):7036–7047.

31. Agamaite JA, Chang C-J, Osmanski MS, Wang X (2015) A quantitative acoustic analysis of the vocal repertoire of the common marmoset (Callithrix jacchus). J Acoust Soc Am 138(5):2906.

32. Epple G (1968) Comparative studies on vocalization in marmoset monkeys (Hapalidae). Folia Primatol Int J Primatol 8(1):1–40.

33. Bezerra BM, Souto A (2008) Structure and usage of the vocal repertoire of Callithrix jacchus. Int J Primatol 29(3):671–701.

34. Roy S, Wang X (2012) Wireless multi-channel single unit recording in freely moving and vocalizing primates. J Neurosci Methods 203(1):28–40.

35. Jasper H, Penfield W (1949) Electrocorticograms in man: Effect of voluntary movement upon the electrical activity of the precentral gyrus. Arch Psychiatr Nervenkr 183(1–2):163–174.

36. Sanes JN, Donoghue JP (1993) Oscillations in local field potentials of the primate motor cortex during voluntary movement. Proc Natl Acad Sci U S A 90(10):4470–4.

37. Murthy VN, Fetz EE (1992) Coherent 25-to 35-Hz oscillations in the sensorimotor cortex of awake behaving monkeys. Proc Natl Acad Sci U S A 89(12):5670–4.

38. Brovelli A, et al. (2004) Beta oscillations in a large-scale sensorimotor cortical network: directional influences revealed by Granger causality. Proc Natl Acad Sci U S A 101(26):9849–54.

39. Baker SN (2007) Oscillatory interactions between sensorimotor cortex and the periphery. Curr Opin Neurobiol 17(6):649–655.

40. Donoghue JP, Sanes JN, Hatsopoulos NG, Gaál G (1998) Neural discharge and local field potential oscillations in primate motor cortex during voluntary movements. J Neurophysiol 79(1):159–173.

41. Engel AK, Fries P (2010) Beta-band oscillations-signalling the status quo? Curr Opin Neurobiol 20(2):156–165.

42. Wessel JR (2020) β-bursts reveal the trial-to-trial dynamics of movement initiation and cancellation. J Neurosci 40(2):411–423.

43. Zaepffel M, Trachel R, Kilavik BE, Brochier T (2013) Modulations of EEG Beta Power during Planning and Execution of Grasping Movements. PLoS One 8(3):e60060.

44. Stančák A, Riml A, Pfurtscheller G (1997) The effects of external load on movement-related changes of the senorimotor EEG rhythms. Electroencephalogr Clin Neurophysiol 102(6):495–504.

45. Spinks RL, Kraskov A, Brochier T, Umilta MA, Lemon RN (2008) Selectivity for grasp in local field potential and single neuron activity recorded simultaneously from M1 and F5 in the awake macaque monkey. J Neurosci 28(43):10961–10971.

46. Stancák A, Pfurtscheller G (1996) Event-related desynchronisation of central beta-rhythms during brisk and slow self-paced finger movements of dominant and nondominant hand. Cogn Brain Res 4(3):171–183.

47. Kilavik BE, et al. (2012) Context-Related Frequency Modulations of Macaque Motor Cortical LFP Beta Oscillations. Cereb Cortex 22(9):2148–2159.

48. Tzagarakis C, Ince NF, Leuthold AC, Pellizzer G (2010) Beta-band activity during motor planning reflects response uncertainty. J Neurosci 30(34):11270–11277.

49. Sochůrková D, Rektor I, Jurák P, Stančák A (2006) Intracerebral recording of cortical activity related to self-paced voluntary movements: A Bereitschaftspotential and event-related desynchronization/synchronization. SEEG study. Exp Brain Res 173(4):637–649.

50. Coupé C, Oh Y, Dediu D, Pellegrino F (2019) Different languages, similar encoding efficiency: Comparable information rates across the human communicative niche. Sci Adv 5(9):eaaw2594.

51. Giraud AL, et al. (2007) Endogenous Cortical Rhythms Determine Cerebral Specialization for Speech Perception and Production. Neuron 56(6):1127–1134.

52. Assaneo MF, Poeppel D (2018) The coupling between auditory and motor cortices is rate-restricted: Evidence for an intrinsic speech-motor rhythm. Sci Adv 4(2):eaao3842.

53. Kingyon J, et al. (2015) High-gamma band fronto-temporal coherence as a measure of functional connectivity in speech motor control. Neuroscience 305:15–25.

54. Behroozmand R, Ibrahim N, Korzyukov O, Robin DA, Larson CR (2015) Functional role of delta and theta band oscillations for auditory feedback processing during vocal pitch motor control. Front Neurosci 9(MAR):109.

55. Gross J, et al. (2002) The neural basis of intermittent motor control in humans. Proc Natl Acad Sci U S A 99(4):2299–2302.

56. Lemke SM, Ramanathan DS, Guo L, Won SJ, Ganguly K (2019) Emergent modular neural control drives coordinated motor actions. Nat Neurosci 22(7):1122–1131.

57. Risueno-Segovia C, Hage SR (2020) Theta Synchronization of Phonatory and Articulatory Systems in Marmoset Monkey Vocal Production. Curr Biol 30(21):4276–4283.e3.

58. Graziano MS., Taylor CS., Moore T (2002) Complex Movements Evoked by Microstimulation of Precentral Cortex. Neuron 34(5):841–851.

59. Wise SP (1985) The primate premotor cortex: past, present, and preparatory. Annu Rev Neurosci 8(1):1–19.

60. Ohbayashi M, Picard N, Strick PL (2016) Inactivation of the Dorsal Premotor Area Disrupts Internally Generated, But Not Visually Guided, Sequential Movements. J Neurosci 36(6):1971–6.

61. Warden MR, Miller EK (2010) Task-dependent changes in short-term memory in the prefrontal cortex. J Neurosci 30(47):15801–10.

62. Hickok G (2012) Computational neuroanatomy of speech production. Nat Rev Neurosci 13(2):135–45.

63. Jurgens U, Ploog D (1970) Cerebral representation of vocalization in the squirrel monkey. Exp brain Res 10(5):532–54.

64. Sutton D, Larson C, Lindeman RC (1974) Neocortical and limbic lesion effects on primate phonation. Brain Res 71(1):61–75.

65. Hage SR, Jürgens U (2006) Localization of a vocal pattern generator in the pontine brainstem of the squirrel monkey. Eur J Neurosci 23(3):840–844.

66. Hage SR, Jürgens U (2006) On the role of the pontine brainstem in vocal pattern generation: a telemetric single-unit recording study in the squirrel monkey. J Neurosci 26(26):7105–15.

67. Düsterhöft F, Häusler U, Jürgens U (2004) Neuronal activity in the periaqueductal gray and bordering structures during vocal communication in the squirrel monkey. Neuroscience 123(1):53–60.

68. Barlow SM, Estep M (2006) Central pattern generation and the motor infrastructure for suck, respiration, and speech. J Commun Disord 39(5):366–380.

69. Dujardin E, Jürgens U (2006) Call type-specific differences in vocalization-related afferents to the periaqueductal gray of squirrel monkeys (Saimiri sciureus). Behav Brain Res 168(1):23–36.

70. Jürgens U, Zwirner P (1996) The role of the periaqueductal grey in limbic and neocortical vocal fold control. Neuroreport 7(18):2921–3.

71. Cerkevich CM, Strick PL (2018) Cortical adaptations to enable enhanced vocalization. Society for Neuroscience Abstract.

72. Lu T, Liang L, Wang X (2001) Neural representations of temporally asymmetric stimuli in the auditory cortex of awake primates. J Neurophysiol 85(6):2364–80.

73. Paxinos G, Watson C, Petrides M, Rosa M, Hironobu T (2012) The Marmoset Brain in Stereotaxic Coordinates (Academic Press).

74. Roy S (2012) Control of vocal production and its neural representation in the primate premotor cortex during natural behavior. PhD Diss.

75. Oostenveld R, Fries P, Maris E, Schoffelen J-M (2011) FieldTrip: Open source software for advanced analysis of MEG, EEG, and invasive electrophysiological data. Comput Intell Neurosci 2011:156869.

76. Tallon-Baudry C, Bertrand O, Delpuech C, Permier J (1997) Oscillatory gamma-band (30-70 Hz) activity induced by a visual search task in humans. J Neurosci 17(2):722–34.

77. Tsunada J, Baker AE, Christison-Lagay KL, Davis SJ, Cohen YE (2011) Modulation of Cross-Frequency Coupling by Novel and Repeated Stimuli in the Primate Ventrolateral Prefrontal Cortex. Front Psychol 2:217.

78. Lu T, Wang X (2000) Temporal Discharge Patterns Evoked by Rapid Sequences of Wide- and Narrowband Clicks in the Primary Auditory Cortex of Cat. J Neurophysiol 84(1):236–246.

79. Eliades SJ, Wang X (2008) Chronic multi-electrode neural recording in free-roaming monkeys. J Neurosci Methods 172(2):201–14.

80. DiMattina C, Wang X (2006) Virtual vocalization stimuli for investigating neural representations of species-specific vocalizations. J Neurophysiol 95(2):1244–62.

81. Eliades SJ, Wang X (2012) Neural correlates of the lombard effect in primate auditory cortex. J Neurosci 32(31):10737–48.

82. Budney GF, Grotke RW (1997) Techniques for Audio Recording Vocalizations of Tropical Birds. Ornithol Monogr (48):147–163.

83. Wahlström S (1985) The Parabolic Reflector as an Acoustical Amplifier. J Audio Eng Soc 33(6):418–429.

